# Biophysical properties of AKAP95 protein condensates regulate splicing and tumorigenesis

**DOI:** 10.1101/536839

**Authors:** Wei Li, Jing Hu, Bi Shi, Francesco Palomba, Michelle A. Digman, Enrico Gratton, Hao Jiang

**Affiliations:** Department of Biochemistry and Molecular Genetics, University of Alabama at Birmingham School of Medicine, Birmingham, AL, USA; Department of Biochemistry and Molecular Genetics, University of Virginia School of Medicine, Charlottesville, VA, USA; Laboratory of Fluorescence Dynamics, The Henry Samueli School of Engineering, University of California, Irvine, CA, USA

## Abstract

It remains unknown if biophysical or material properties of biomolecular condensates regulate cancer. Here we show that AKAP95, a nuclear protein that regulates transcription and RNA splicing, plays an important role in tumorigenesis by supporting cancer cell growth and suppressing oncogene-induced senescence. AKAP95 forms phase-separated and liquid-like condensates in vitro and in nucleus. Mutations of key residues to different amino acids perturb AKAP95 condensation in opposite directions. Importantly, the activity of AKAP95 in splice regulation is abolished by disruption of condensation, significantly impaired by hardening of condensates, and regained by substituting its condensation-mediating region with other condensation-mediating regions from irrelevant proteins. Moreover, the abilities of AKAP95 in regulating gene expression and supporting tumorigenesis require AKAP95 to form condensates with proper liquidity and dynamicity. These results link phase separation to tumorigenesis and uncover an important role of appropriate biophysical properties of protein condensates in gene regulation and cancer.

## INTRODUCTION

Emerging as a fundamental principle in organizing cellular contents and biochemistry, phase separation or biomolecular condensation is driven by multivalent and weak interactions among nucleic acids and proteins that often involve intrinsically disordered regions (IDRs)^1–5^. Biomolecular condensates can adopt a broad spectrum of material properties, from highly dynamic liquid to semi-fluid gels and solid amyloid aggregates^4, 6–8^. Pathogenic mutations can facilitate high-degree aggregation that underlies certain degenerative diseases^7, 9–12^. However, we don’t understand well the role of phase separation in other major diseases, such as cancer, which arises from genetic alterations that often reprogram gene transcription^13^ and sometimes RNA splicing^14^. Phase separation is linked to spatiotemporal regulation of gene expression^15^. Transcription involves condensation of proteins including RNA polymerase II (Pol II)^16, 17^, transcription factors^18, 19^, and coactivators^20^. Aggregation or condensation of splicing regulators mediates RNA splice activation^21^ and may expand their gene regulatory capacity in mammals^22^. The condensation property of FUS, EWSR1, and TAF15 is implicated in the oncogenic potentials of their translocated products through transcriptional regulation^16, 23^, and the condensation domain of the EWS-FLIi fusion recruits chromatin-remodelers to drive oncogenic gene expression^24^. However, it remains unknown whether different liquidity and dynamics functionally impact biochemical outcomes in gene expression and biological consequences in cancer.

AKAP95 (also called AKAP8)^25^ is a nuclear member of the A-kinase anchoring proteins family with multiple activities^26–30^, and also a member of the AKAP95 family with the common subtype of the zinc finger (ZF) domains^31^. We have previously shown that AKAP95 integrates regulation of transcription and RNA splicing^32, 33^. Through its 1-100 region, AKAP95 binds many factors in RNA processing and transcription, including DDX5, a subset of hnRNPs, and Pol II^33^. AKAP95 co-activates expression of a chromatin reporter^32^, directly regulates splicing of a minigene reporter, and modulates alternative splicing of human transcriptome by directly binding to pre-mRNA introns in a ZF-dependent manner^33^. However, the fundamental molecular properties underlying its activity in gene regulation are unclear. The pathophysiologic role of AKAP95 is also poorly understood, other than its implications in diseases including abnormal head growth, autism^34^, and prenatal oral clefts^35^. Consistent with enrichment of cell cycle-related transcripts in AKAP95 targets^33^, AKAP95 is overexpressed in clinical samples of primary ovarian^36^ and rectal cancer tissues^37^ together with some cyclins. We thus set out to determine the fundamental molecular properties of AKAP95 in regulating gene expression and cancer. Our results show that AKAP95 plays an important role in supporting tumorigenesis through splice regulation, which requires AKAP95 to form protein condensates with proper dynamics.

## RESULTS

### AKAP95 is associated with human cancers and regulates cancer cell growth

Our analyses show that *AKAP95* is frequently overexpressed across a large variety of human cancers (**Extended Data Fig. 1a**), and its amplification and upregulation in triple negative breast cancer (TNBC) are correlated with poorer patient survival (**Fig. 1a**). AKAP95 knockdown (KD) in the MDA-MB-231 TNBC cells markedly reduced growth in culture, with reduced proliferation and increased apoptosis, and reduced in vivo tumorigenicity (**Fig. 1b,c, Extended Data Fig. 1b**). Restoring AKAP95 expression rescued cell growth, and extra expression of AKAP95 in control cells enhanced growth (**Extended Data Fig. 1c**). The requirement of AKAP95 in breast cancer cell growth is not limited to TNBC, as shown in the estrogen receptors-positive MCF-7 cells (**Extended Data Fig. 1d**). RNA-seq analysis (**Supplementary Table 1, tab 1)** showed that the TGF-β signaling, G2M checkpoint, and E2F targets genes were significantly downregulated, and the inflammatory response and apoptosis genes were upregulated by KD (**Fig. 1d,e, Extended Data Fig. 1e**). Moreover, AKAP95 KD changed thousands of alternative splicing events enriched in pathways including DNA damage and repair, cell division, mitochondrion, and transcription (**Fig. 1f,g, Supplementary Table 1, tab 2**).

**Fig. 1.**
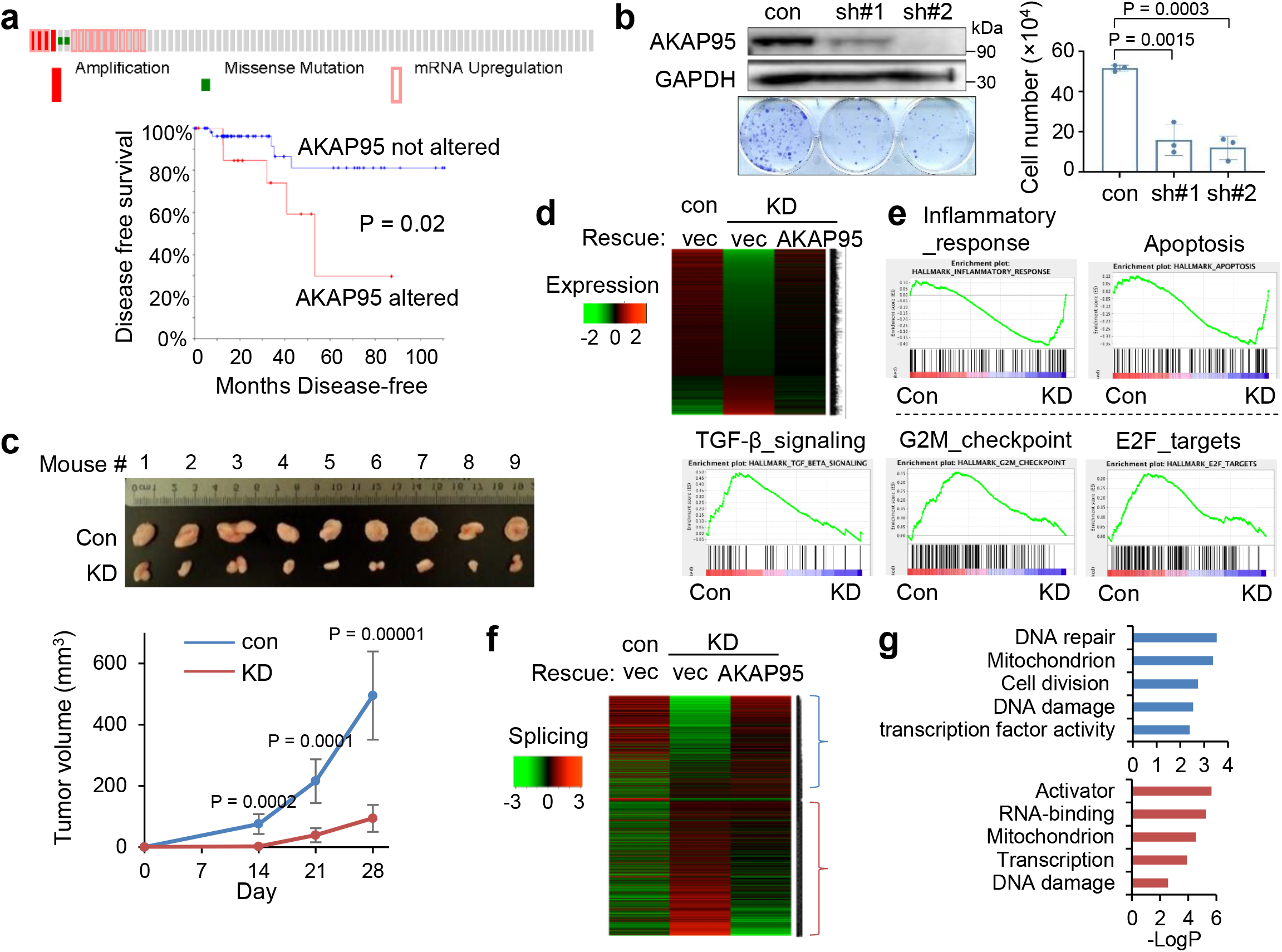
AKAP95 regulates cancer cell growth and gene expression. **a,** Overexpression of *AKAP95* in breast cancer tissues of 82 TNBC patient samples. From cBioPortal. Top, each box is a patient sample. Bottom, disease-free survival curves of patients with or without *AKAP95* alterations. **b,** Growth assay for MDA-MB-231 cells expressing control or two AKAP95 shRNAs. Left, immunoblotting of total cell lysates and images of cell colonies stained with crystal violet. Right, numbers of cells in growth assays as mean ± SD from n = 3 independent experiments. **c,** Tumors from xenograft of control or AKAP95-KD MDA-MB-231 cells in immune-deficient mice. Tumor volumes at the indicated days post transplantation are plotted as mean ± SD (n = 9). **d,e,** RNA-seq analysis in MDA-MB-231 cells expressing control or AKAP95 shRNA #1 and the indicated vector or AKAP95-expressing construct. **d,** Heatmap showing relative expression levels of genes down- or up-regulated in the indicated cells. It includes 951 and 294 genes down- and up-regulated in KD compared to control cells, respectively. Also see Supplementary Table 1, tab 1. **e,** GSEA for gene expression profiles of control and AKAP95-KD cells. Plots above and below the broken line show gene sets significantly enriched in up- and down-regulated genes by AKAP95 KD, respectively. **f,** Heatmap showing relative alternative splicing and clustered by changes in percent-spliced-in (PSI) values in the indicated cells. It includes 807 and 1275 alternative splicing events with decreased or increased PSI in KD cells, respectively. Also see Supplementary Table 1, tab 2. **g,** Gene ontology analysis for the indicated clusters from the heatmap in f. Blue (n = 807) and red (n = 1275) show functions significantly enriched in genes with PSI increase or decrease by AKAP95 KD, respectively. P values by log-rank test for a, Student’s *t*-test for b and d, and modified Fisher’s exact test for g. All two-sided. Uncropped blots are provided as source data.

### AKAP95 directly regulates splicing of cancer-related targets

Consistent with the effects on proliferation, AKAP95 KD selectively downregulated *CCNA2* expression (**Fig. 2a**). *CCNA2* and *AKAP95* are often co-overexpressed in TNBC patients (**Fig. 2b**). CCNA2 KD inhibited growth of TNBC cells, and its restoration in the AKAP95-KD cells rescued the growth (**Extended Data Fig. 2a,b**). AKAP95 binds to intron 1 of *CCNA2* pre-mRNA in a ZF-dependent manner (**Fig. 2c**), and AKAP95 KD reduced the relative junction level of exons 1 and 2 (**Extended Data Fig. 2c**). As *CCNA2* intron 1 harbors a premature translation termination codon known to activate nonsense-mediated decay (NMD)^38^, we detected retention of *CCNA2* intron 1 following AKAP95 KD when NMD was blocked by either cycloheximide or depletion of UPF1 or BTZ, key proteins in NMD^38^ (**Fig. 2d, Extended Data Fig. 2c-e**). Moreover, NMD blockage significantly stabilized *CCNA2* mRNA in AKAP95-KD cells, but not in control cells (**Fig. 2e**). Therefore, AKAP95 directly facilitates *CCNA2* splicing to ensure its sufficient production that contributes to cancer cell growth.

**Fig. 2.**
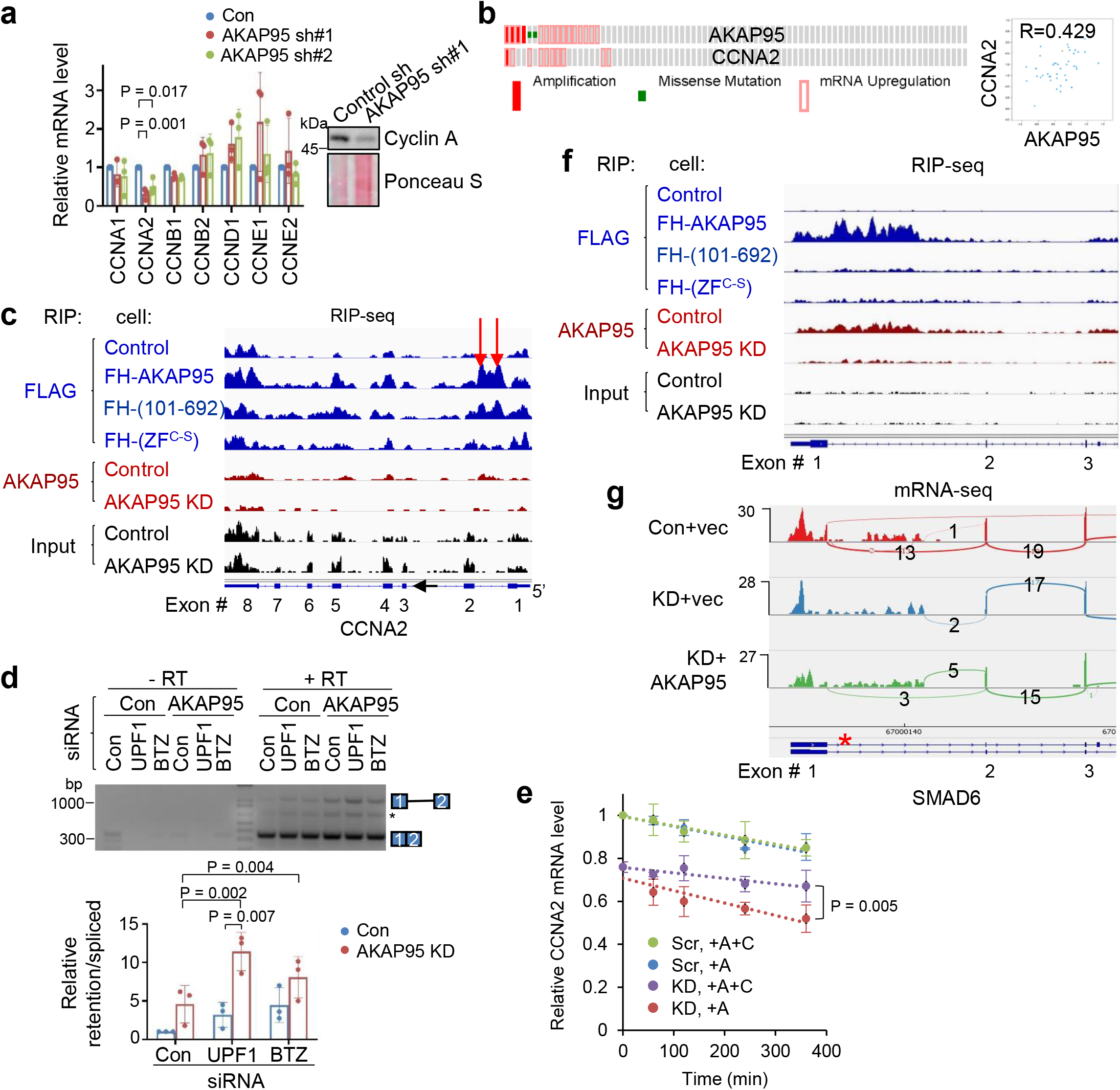
AKAP95 directly regulates splicing of key transcripts for cancer. **a,** *CCNA2* expression in MDA-MB-231 cells upon AKAP95 KD. Left, relative mRNA levels of indicated cyclins were determined by RT-qPCR and normalized to *GAPDH*, and presented as mean ± SD from n = 3 biological repeats. Right, immunoblotting for Cyclin A1/A2 in total cell lysates. **b,** Co-overexpression of *AKAP95* and *CCNA2* in breast cancer tissues of 82 TNBC patients. Left, each box represents a patient. Right, correlation of their mRNA levels in the TNBC patients with indicated Pearson correlation coefficient. Both from cBioPortal. **c,f,** RNA immunoprecipitation-sequencing (RIP-seq) profiles for *CCNA2* (c) and *SMAD6* (f) based on our previous work^33^. In blue are Anti-FLAG RIP-seq in control or 293 cells expressing the FLAG-HA-tagged AKAP95 WT or mutants. In red are anti-AKAP95 RIP-seq in control or AKAP95-KD 293 cells. In black are profiles of total input RNAs. All profiles have the same Y-axis scale. Red arrows indicate AKAP95-binding sites at intron 1. **d,** Total RNAs were used for RT-PCR, in the absence (-RT) or presence (+ RT) of reverse transcriptase, for *CCNA2* intron 1 in MDA-MB-231 cells with indicated combination of siRNAs. Top, PCR products on agarose gel. Asterisk, an unknown amplification product. Repeated 3 times. Bottom, relative ratios of the signal for the intron 1-retaining transcript over the intron 1-spliced transcript, as mean ± SD from n = 3 independent experiments. **e,** Assay for *CCNA2* mRNA stability. Control (Scr) and AKAP95-KD MDA-MB-231 cells were treated starting from 0 min with Actinomycin D (+A, to block RNA synthesis) and cycloheximide (+C, to block NMD) or not as indicated. Total RNA at indicated times were used for RT-PCR and normalized to *ACTB*, as mean ± SD from n = 3 biological repeats. **g,** mRNA-seq profiles for *SMAD6* in MDA-MB-231 cells expressing control or AKAP95 shRNA #1 (KD) and vector or AKAP95-expressing construct. Asterisk, a stop codon. P values by two-sided Student’s *t*-test for a and e and one-way ANOVA followed by Tukey’s post hoc test for d. Uncropped blots are provided as source data.

SMAD6^39^, a negative regulator of TGF-β signaling^40^ known to promote tumorigenesis^41, 42^, was among this signaling pathway genes downregulated upon AKAP95 KD and rescued by AKAP95 restoration (**Extended Data Fig. 2f**). AKAP95 bound to intron 1 of *SMAD6* pre-mRNA (**Fig. 2f**). AKAP95 KD abolished joining of exons 1-2 but not 2-3, and thus may activate NMD of the intron 1-retained *SMAD6* transcript containing a translation termination codon (**Fig. 2g**). AKAP95 also substantially bound to intronic regions flanking an AKAP95-controlled alternative exon of two other exemplary gene transcripts *RPUSD3* and *PPM1K* (**Fig. 8f, Supplementary Figs. 2g,h,8c**). Taken together, these results suggest that AKAP95 regulates cancer cell growth partially through controlling the splicing and expression of cancer-related targets.

### AKAP95 is dispensable for normal cell growth but required for suppressing oncogene-induced senescence

To better understand the role of AKAP95 in cancer, we generated an *Akap95* knockout (KO) mouse model (**Extended Data Fig. 3a**). *Akap95^−/−^* mice were born with expected Mendelian ratio, and had no overt phenotypes, including largely unaffected body weight and blood cell profiling (**Extended Data Fig. 3b,c**). AKAP95 is thus dispensable for normal animal development or physiology. We derived primary mouse embryonic fibroblasts (MEFs) from wild type (WT), *Akap95^+/−^* (Het), and *Akap95^−/−^* (KO) embryos (**Extended Data Fig. 3d**). The Het and KO MEFs showed no difference in growth rate (**Fig. 3a**) or morphology (**Figs. 3e**). However, upon expression of two potent oncogenes, *H-RAS^G12V^* and c-*MYC* (*MYC*) at levels unaffected by AKAP95 loss, the KO MEFs showed reduced anchorage-independent colony formation and in vivo tumorigenicity (**Fig. 3b-3d**). Therefore, while dispensable for normal cell growth, AKAP95 is required for oncogene-induced transformation.

**Fig. 3.**
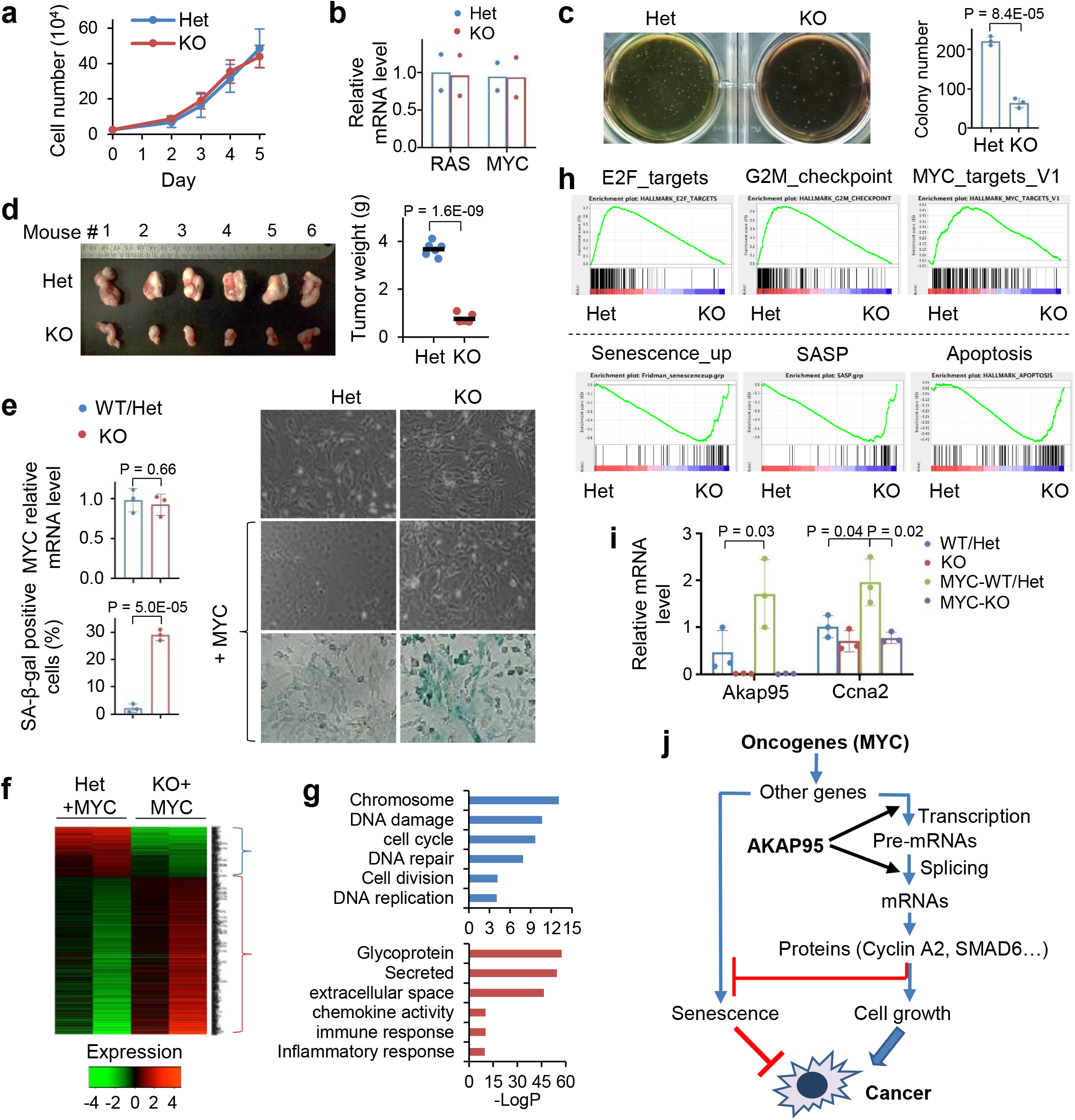
AKAP95 is dispensable for normal cell growth but required for transformation and suppressing oncogene-induced cellular senescence. **a,** Growth of MEFs from *Akap95*^+/−^ (Het) and *Akap95*^−/−^ (KO) embryos (n = 6 each). **b,** Relative *HRAS* and *MYC* mRNA levels were determined by RT-qPCR and normalized to *Actb*, as mean ± SD from HRAS^G12V^ and MYC transduced MEFs of 2 embryos each. **c,** HRAS-MYC-transduced MEFs in soft agar colony formation assay. Colony numbers as mean ± SD from n = 6 experiments using MEFs of 2 embryos each. **d,** Six mice received HRAS-MYC-transduced Het and KO MEFs on each flank. Tumor weights at four weeks are plotted. Each dot represents a tumor. **e-i,** MEFs derived from 3 KO and 3 Akap95-expressing (containing 1 WT and 2 Het) embryos were transduced with MYC. **e,** Right, images of cells before and after MYC transduction. Images of SA-beta-galactosidase activity assay are at the bottom. Relative *MYC* mRNA levels after transduction were determined by RT-qPCR and normalized to *Actb* (left top). Percentage of SA-beta-gal-positive cells are plotted (left bottom). Both as mean ± SD from MEFs (n = 3 embryos each). **f,** Heatmap showing relative expression levels of genes and clustered by changes in KO MEFs (2 embryos each). It includes 265 and 742 genes down- or up-regulated in KO, respectively. Also see Supplementary Table 2, tab 1. **g,** Gene ontology analysis for the indicated gene clusters from the heatmap in f. Blue (n = 265) and red (n = 742) show functions significantly enriched in down- and up-regulated genes, respectively. **h,** GSEA plots above and below the dashed line show gene sets significantly enriched in genes down- and up-regulated in the MYC-transduced KO compared to Het MEFs, respectively. **i,** Relative *Akap95* and *Ccna2* mRNA levels before and after MYC transduction as determined by RT-qPCR and normalized to *Actb*, as mean ± SD from MEFs from 3 KO and 2 WT/Het embryos. **j,** A diagram summarizing regulation of tumorigenesis by AKAP95 through gene expression control. P values by two-sided Student’s *t*-test for all except one-way ANOVA followed by Tukey’s post hoc test for i, and modified Fisher's exact test for g. Uncropped blots are provided as source data.

Upon transduction of MYC alone, the KO MEFs did not exhibit substantial morphological changes seen for the WT MEFs (as expected for cells being immortalized), and showed higher levels of senescence marker than WT (**Fig. 3e**). In the absence of MYC transduction, AKAP95 loss altered (mostly downregulated) expression of a limited number of genes (**Extended Data Fig. 3e, Supplementary Table 2, tab 6**). In the MYC-transduced MEFs, however, AKAP95 loss altered (mostly upregulated) expression of a much larger number of genes, including downregulation of nuclear pathways in cell proliferation and DNA damage repair, and upregulation of senescence, apoptosis, and extracellular pathways in inflammatory response, especially the senescence-associated secretory phenotype (SASP) that helps eliminate malignant cells undergoing senescence^43, 44^ (**Figs. 3f-h, Extended Data Fig. 3f, Supplementary Table 2, tab 1**). The global gene expression alteration could be largely reverted by introduction of WT AKAP95 (**Extended Data Fig. 3g, Supplementary Table 2, tab 2**). These results indicate that AKAP95 is required for removing the senescence barrier on the path to cancer.

MYC transduction in the Het MEFs markedly upregulated the expression of *Akap95*, but not its close paralog, *Akap8l* (**Extended Data Fig. 3h**). Moreover, MYC upregulated *Ccna2* in the Het but not KO MEFs (**Fig. 3i**). Moreover, AKAP95 loss altered hundreds of events in alternative exon inclusion (**Extended Data Fig. 3e,i,k, Supplementary Table 2, tabs 3, 7**). Similar to the TNBC cells, the KO-affected genes in MYC-transduced MEFs were also enriched in pathways including DNA damage and repair, cell division, mitochondrion, and transcription (**Extended Data Fig. 3j**). Therefore, in oncogenic assault (e.g., MYC), the activity of AKAP95 in regulating gene expression becomes important for tumorigenesis by supporting cell proliferation and suppressing stress response such as senescence (**Fig. 3j**).

### AKAP95 forms liquid-like and phase-separated condensates in vitro

Our observations of AKAP95 forming dimers resistant to strongly denaturing conditions (**Fig. 4a, Extended Data Fig. 4a)** suggested its extraordinary propensity in self-aggregation. AKAP95 is predicted to be intrinsically disordered throughout the protein except 1-100 and ZFs (**Fig. 4b**). We purified different AKAP95 regions individually fused to maltose-binding protein (MBP, to enhance yield and solubility) (**Extended Data Fig. 4c**). AKAP95 (1-340), but not (306-692), exhibited remarkable turbidity with elevated OD600 upon cleavage of MBP (**Fig. 4c, Extended Data Fig. 4d**). Consistent with 101-210 mediating AKAP95 self-interaction^33^, this region was sufficient and necessary to develop turbidity (**Fig. 4c, Supplementary Figs. 4d**),

**Fig. 4.**
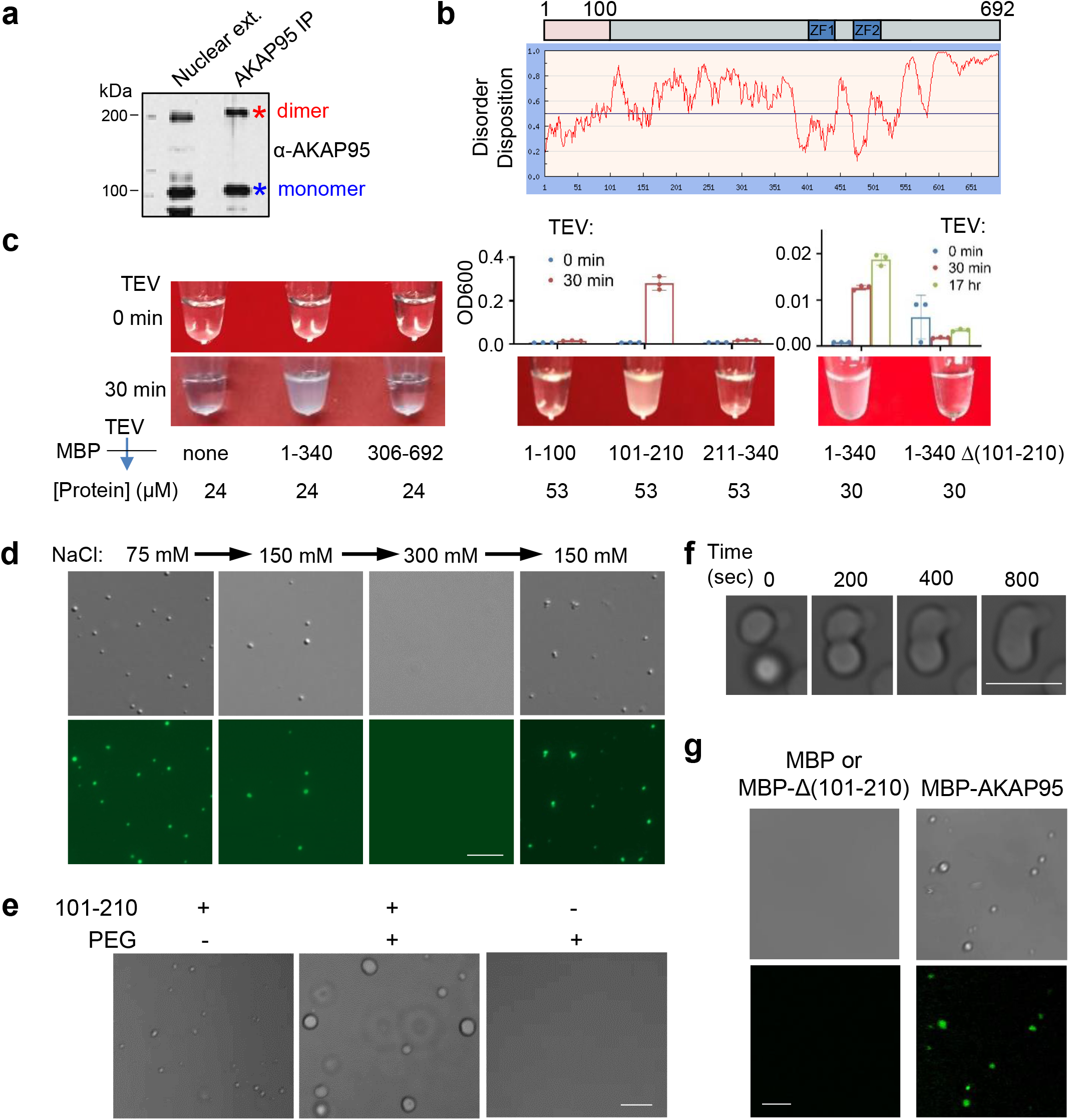
AKAP95 undergoes phase separation with liquid-like properties in vitro. **a,** Immunoblotting for AKAP95 in HeLa cell nuclear extract and AKAP95 immunoprecipitation from the extract. Samples were boiled in the presence of DTT and resolved by SDS-PAGE. **b,** Disorder plot of human AKAP95. **c,** Turbidity by pictures and OD600 of MBP (none) and MBP fused to AKAP95 truncations at indicated concentrations all in 30 mM NaCl before and after TEV protease treatment for indicated time. OD600 is plotted as mean ± SD from n = 3 biological repeats. **d,** DIC (top) and fluorescence microscopy (bottom) images for 20 μM MBP-AKAP95 (101-210) and spiked with Oregon-green-labeled same protein (molar ratio 10:1) after TEV protease treatment for 30 min. Changes in NaCl concentration is indicated. Images were taken 5 min after salt adjustment. **e,** Phase contrast images of 50 μM MBP-AKAP95 (101-210) in 30 mM NaCl in the absence and presence of 10% of PEG6000 after TEV protease treatment for 30 min. **f,** Fusion of two droplets formed by 50 μM MBP-AKAP95 (101-210) in 30 mM NaCl and 10% of PEG6000 after TEV protease treatment for 30 min. Also see Movie 1. **g,** DIC and fluorescence microscopy images of 6.25 μM MBP, MBP fused to Δ(101-210) or full-length AKAP95 in 150 mM NaCl, spiked with Oregon-green-labeled AKAP95 (101-210) at a molar ratio of 150:1 after TEV protease treatment for 30 min. Note that the lack of any condensates in the DIC images showed the inability of Δ(101-210) in condensation. Experiments in a, d, e-g were Repeated > 3 times. Scale bar, 5 μm for all. Uncropped blots are provided as source data.

Fluorescently labeled AKAP95 (101-210) formed micron-sized droplets that had nearly spherical shape, freely moved in solution, fell onto glass coverslip, became enlarged in the presence of crowding agent, and occasionally experienced fusion (**Figs. 4d-4f, Extended Data Fig. 4e, Supplementary Movie 1**). The pre-formed droplets were rapidly dissolved by elevated salt concentration and re-formed upon dilution to lower salt (**Fig. 4d**), indicating their high reversibility and dependence on electrostatic interactions. Fluorescence Recovery After Photobleaching (FRAP) assays showed that molecules in the droplets were dynamic (**Fig. 7c**). At a concentration on the same order of magnitude as its endogenous nuclear concentration [2-3 μM in 293 and MDA-MB-231 cell nuclei (**Extended Data Fig. 4f**)], purified full-length AKAP95 also formed condensates in a physiological salt condition in a (101-210)-dependent manner (**Fig. 4g**). These results indicate that AKAP95 undergoes phase separation in vitro and 101-210 confers the liquid-like property of AKAP95 condensates.

**Fig. 7.**
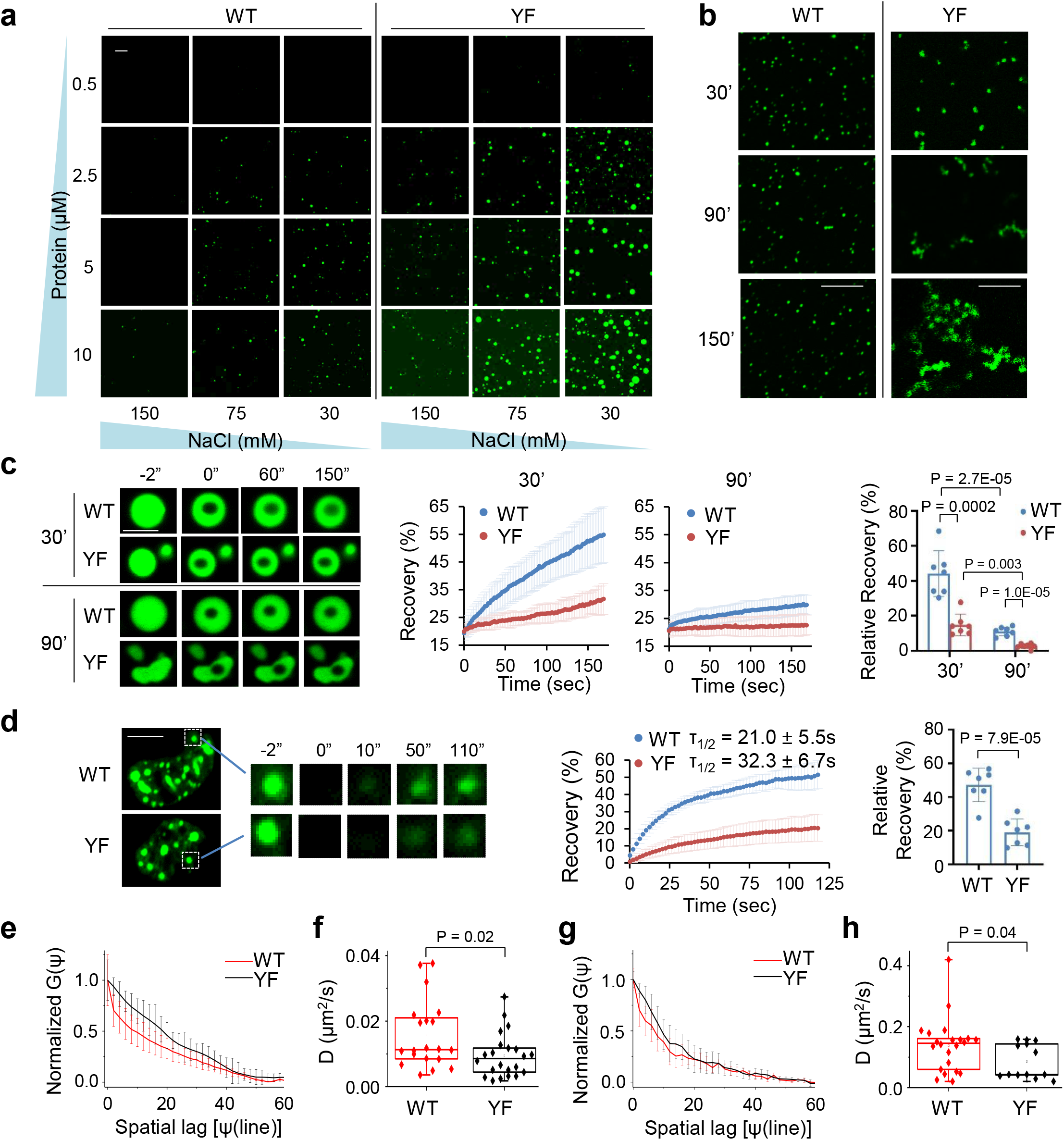
YF mutation enhances AKAP95 condensation propensity and renders condensates more solid-like state. **a,** Fluorescence microscopy images of Oregon-green-labeled AKAP95 (101-210) WT and YF at indicated protein and NaCl concentrations after TEV protease treatment for 30 min. Scale bar, 10 μm. Repeated > 3 times. **b,** Fluorescence microscopy images of 50 μM AKAP95 (101-210) WT and YF in 30 mM NaCl both spiked with Oregon-green-labeled (101-210) WT (molar ratio 150:1) after TEV protease treatment for 30’ and imaged immediately (30’) or after incubation for 60 (90’) or 120 (150’) more minutes. Scale bar, 10 μm. Repeated > 3 times. **c,** FRAP of 10 μM GFP-AKAP95 (101-210) WT and YF after 30 min of MBP cleavage in 150 mM NaCl and 10% of PEG6000. FRAP was performed immediately (30’) or after incubation for 60 min more (90’). Left, fluorescence microscopy images of droplets at indicated times. Middle, FRAP recovery curves as mean ± SD. Right, mean ± SD of recovery (relative to minimal level) at the final time. n = 7 droplets each. Scale bar, 2 μm. **d,** FRAP of Full-length AKAP95 WT and YF fused to GFP in HeLa cell nuclei. Left, fluorescence microscopy images of foci. The photobleached focus was boxed and amplified for the indicated time points. Middle, FRAP recovery curves as mean ± SD. Right, mean ± SD of recovery (relative to minimal level) at the final time. n = 7 cells each. Scale bar, 5 μm. **e,f,** Diffusion of full length AKAP95 WT and YF fused to GFP in HeLa cell nuclei, showing Line RICS normalized autocorrelation curves G(Ψ) as a function of the Spatial Lag (Ψ) (**e**) and diffusion coefficients (**f**), both as mean ± SD (n = 20 for WT, n = 22 for YF). **g,h,** Diffusion coefficients of purified GFP-AKAP95 (101-210) WT and YF, showing Line RICS normalized autocorrelation curves G(Ψ) (**g**) and diffusion coefficients (**h**), both as mean ± SD (n = 22 for WT, n = 13 for YF). P values by two-sided Student’s *t*-test (c, d) or Mann-Whitney *U* test (f, h). For box and whisker plots, data are median (line), 25–75th percentiles (box) and minimum-maximum values recorded (whiskers). Uncropped blots are provided as source data.

**Fig. 8.**
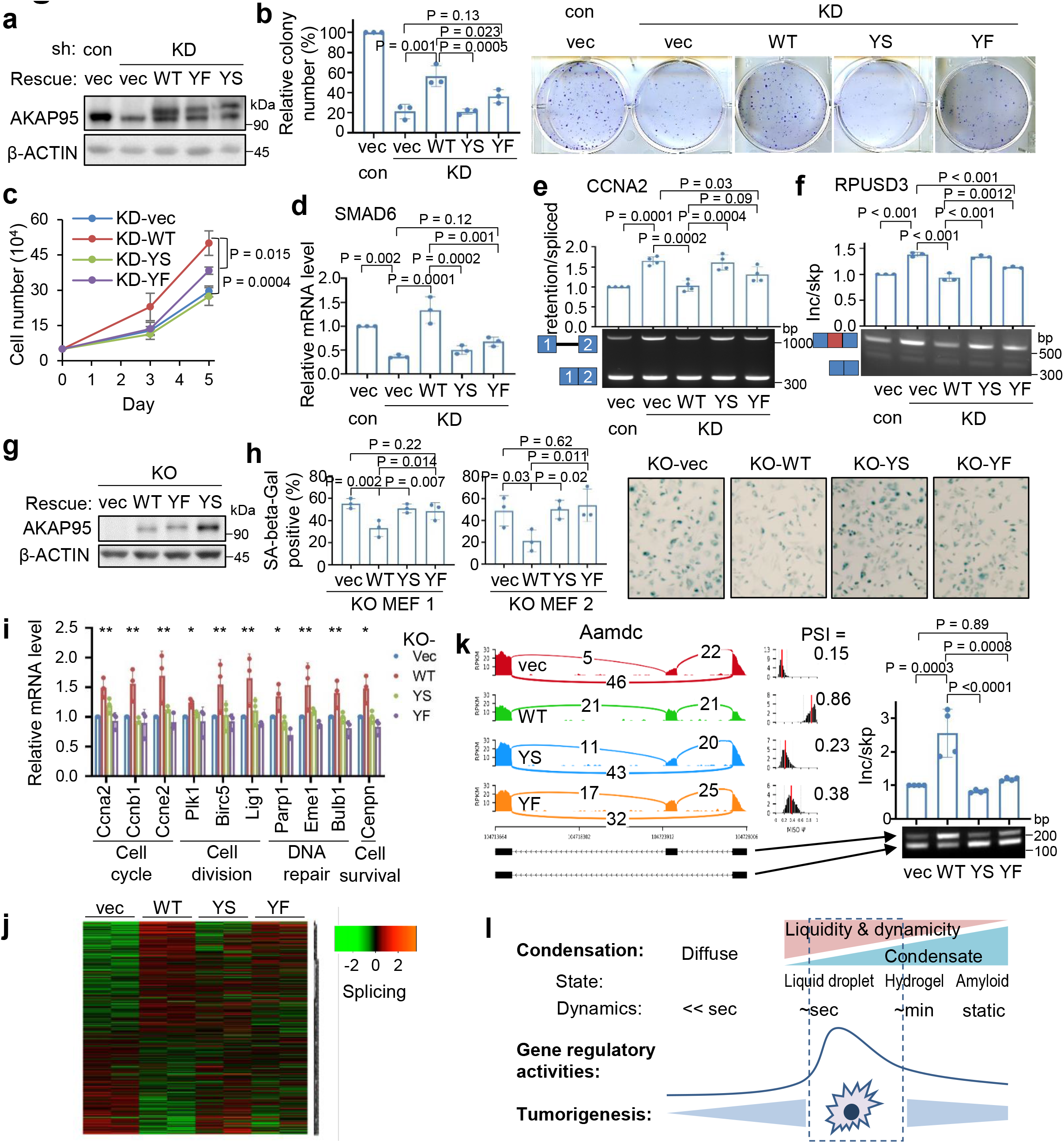
Regulation of tumorigenesis and gene expression by AKAP95 requires its condensation with proper biophysical properties. **a-f,** MDA-MB-231 cells were transduced with control or AKAP95 shRNA #1 (KD) and vector and FLAG-HA-tagged full-length AKAP95 WT or mutants. **a,** Immunoblotting of total cell lysates. Repeated 3 times. **b,** Colony formation assays. Left, colony numbers as mean ± SD from n = 3 biological repeats. Right, images of cells stained with crystal violet. **c,** Growth of cultured cells, as mean ± SD from n = 3 independent experiments. **d,** Relative *SMAD6* mRNA level were determined by RT-qPCR and normalized to *GAPDH*, as mean ± SD from n = 3 biological repeats. **e,** RT-PCR for ratios for intron 1-retained over-spliced *CCNA2* transcript, as mean ± SD from n = 4 biological repeats. **f,** RT-PCR for ratios for exon-included over-skipped *RPUSD3* transcript, as mean ± SD from n = 3 biological repeats. **g-k,** MYC-transduced *Akap95* KO MEFs were transduced with vector or constructs expressing HA-tagged full-length AKAP95 WT or mutants. **g,** Immunoblotting of total cell lysates. Repeated 3 times. **h,** Left, percentage of SA-beta-gal-positive cells as mean ± SD from n = 3 different images of MEFs from two embryos. Right, images from KO MEF 1. **i,** Relative mRNA levels of indicated genes, with related functions at bottom, were determined by RT-qPCR and normalized to *Gapdh*, as mean ± SD from n = 3 independent experiments. * or ** between vec and WT, WT and YS, WT and YF, except for Plk1, for which * only between vec and WT, WT and YF. *P<0.05, **P<0.01. **j,** Heatmap showing relative alternative splicing with PSI changes in MYC-transduced KO MEFs expressing indicated constructs (2 embryos each). Also see Supplementary Table 2, tab 5. **k,** Sashimi plot showing *Aamdc* alternative splicing that was rescued by introduction of AKAP95 WT, but not but the mutant, and RT-PCR for the inclusion of the alternative exon as mean ± SD from n = 4 biological replicates pooled from 2 embryos each. **l,** Diagram showing impact of material properties of AKAP95 WT and mutants on gene regulation and tumorigenesis. P values by two-sided Student’s *t*-test for c and one-way ANOVA followed by Tukey’s post hoc test for all other analyses. Uncropped blots are provided as source data.

### AKAP95 forms dynamic liquid-like droplets in cell nucleus

Endogenous AKAP95 showed punctate distribution in nuclei of human cancer cells and primary MEFs (**Fig. 5a**). Full-length AKAP95, but not the mutant lacking 101-210 [Δ(101-210)], formed nuclear bodies of near spherical shape (**Figs. 5b,c**). These foci increased in size and decreased in number over time (**Fig. 5d**), likely due in part to merging. Interestingly, the AKAP95 (ZF^C-S^) RNA-binding mutant^33^ formed droplets that appeared more spherical and showed rapid fusion in nucleus (rare for WT foci) (**Figs. 5b,e, Supplementary Movie 2**), suggesting that RNA binding to ZFs restrains AKAP95 mobility. A short characteristic recovery time in FRAP assays (**Fig. 7d, Supplementary Movie 3**) indicates that AKAP95 rapidly exchanged in and out of the bodies. These results are consistent with a dynamic liquid-like property of the AKAP95 nuclear foci.

**Fig. 5.**
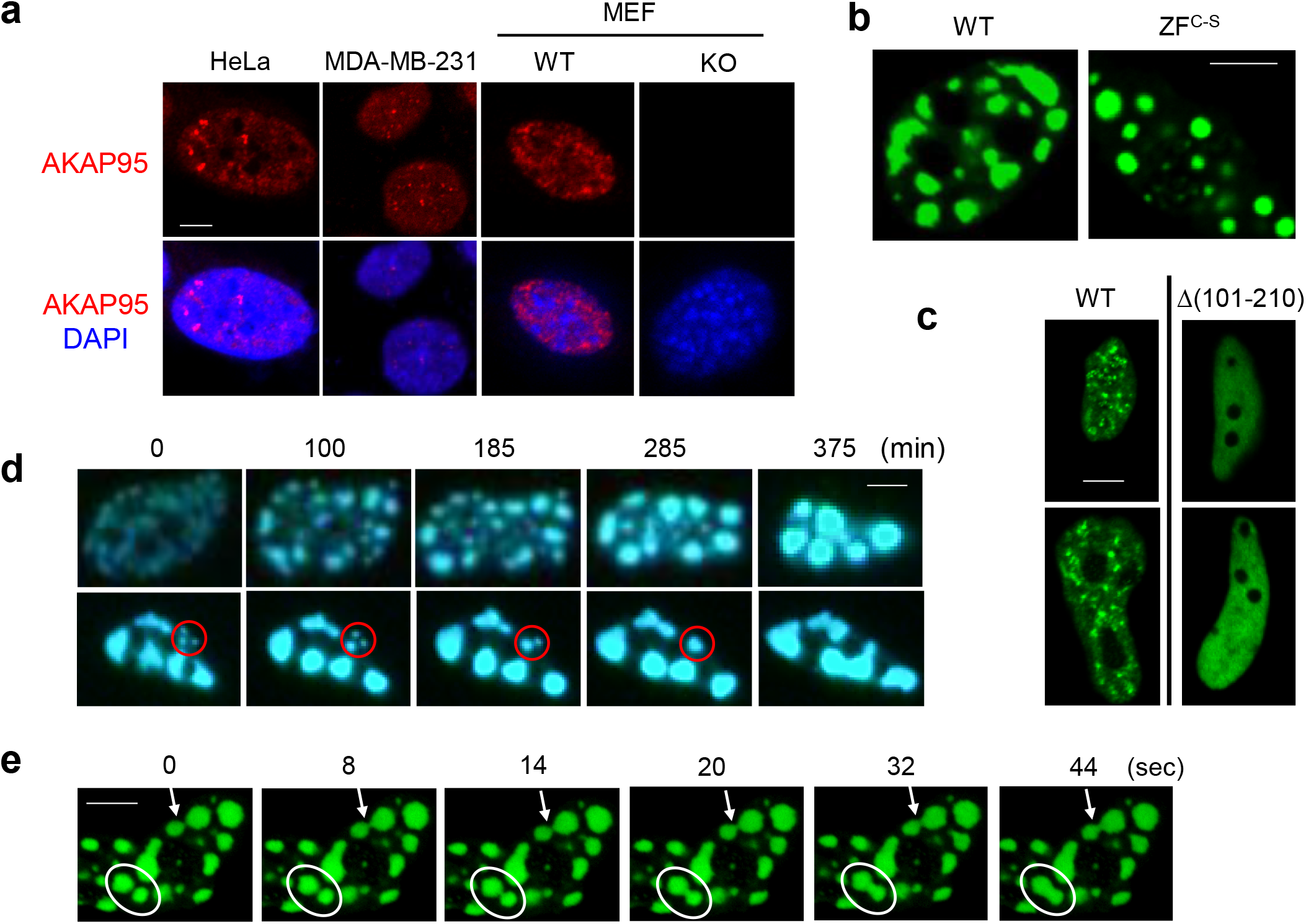
AKAP95 forms dynamic foci in cell nucleus. **a,** Immunostaining of endogenous AKAP95 (red) and DNA (DAPI, blue) in indicated cancer cell lines and primary MEFs from WT and *Akap95* KO embryos. **b,** Confocal microscopy images of AKAP95 WT or ZF^C-S^ fused to GFP in nuclei following transfection into HeLa cells. **c,** Fluorescence microscopy images of HeLa cells transiently expressing AKAP95 WT or Δ(101-210 fused to GFP. **d,** HeLa cells were transfected with AKAP95-GFP, and two nuclei were imaged at different time points. Time 0 was 24 hr after transfection. Note the growth and merge of the foci, especially those in the red circle. **e,** Rapid fusion of AKAP95 (ZF^C-S^)-GFP foci in a HeLa cell nucleus. The white oval and arrow show two different fusion events. These images are from movie 2. All experiments were Repeated > 3 times. Scale bar, 5 μm for all.

AKAP95 is known to associate with chromatin^28, 45^ and nuclear matrix^46^, but its exact location in nucleus is unclear. We found that AKAP95 substantially but incompletely overlapped with SRSF2 (**Extended Data Fig. 5a**), a splicing factor enriched in nuclear speckles^47^. AKAP95 puncta also frequently overlapped with Pol II phosphorylated at Ser 2 of the C-terminal domain, and had little overlap with unphosphorylated Pol II (**Extended Data Fig. 5b,c**). As Ser 2 phosphorylation drives Pol II from the transcriptional condensates to the splicing condensates^48^, these results further support the close involvement of AKAP95 in RNA splicing associated with active transcription.

### Phase separation property is required for AKAP95 to regulate splicing

Consistent with its intrinsic disorder, AKAP95 (101-210) is enriched in flexible amino acids and devoid of bulky residues (**Figs. 6a**, **Extended Data Fig. 6a**). This region is highly conserved between human and mouse, especially for Arg, Asp, Gly, Phe, and Tyr (**Fig. 6a**). Considering the emerging role of Tyr in protein condensation^16, 49, 50^, we mutated all six Tyr residues in 101-210 to Ala (YA), Ser (YS), or Phe (YF) and purified these proteins (**Extended Data Fig. 6b**). Although these mutations were predicted not to substantially alter the disorder status (**Extended Data Fig. 6c**), we found that WT and YF, but not YA and YS, **(i)** appeared turbid with elevated OD600 (YF > WT) (**Fig. 6b**), **(ii)** were markedly depleted from supernatant and enriched in pellet after centrifugation (**Fig. 6b**, **Extended Data Fig. 6d**), and **(iii)** formed micrometer-sized spherical droplets (**Fig. 6c**). Full-length WT or YF, but not YA and YS, either overexpressed at the comparable level or all adjusted to near endogenous level, formed nuclear foci (**Fig. 6d, Extended Data Fig. 6e-g**). The same dependence on Tyr suggests the same molecular nature of assemblies in vitro and in vivo.

**Fig. 6.**
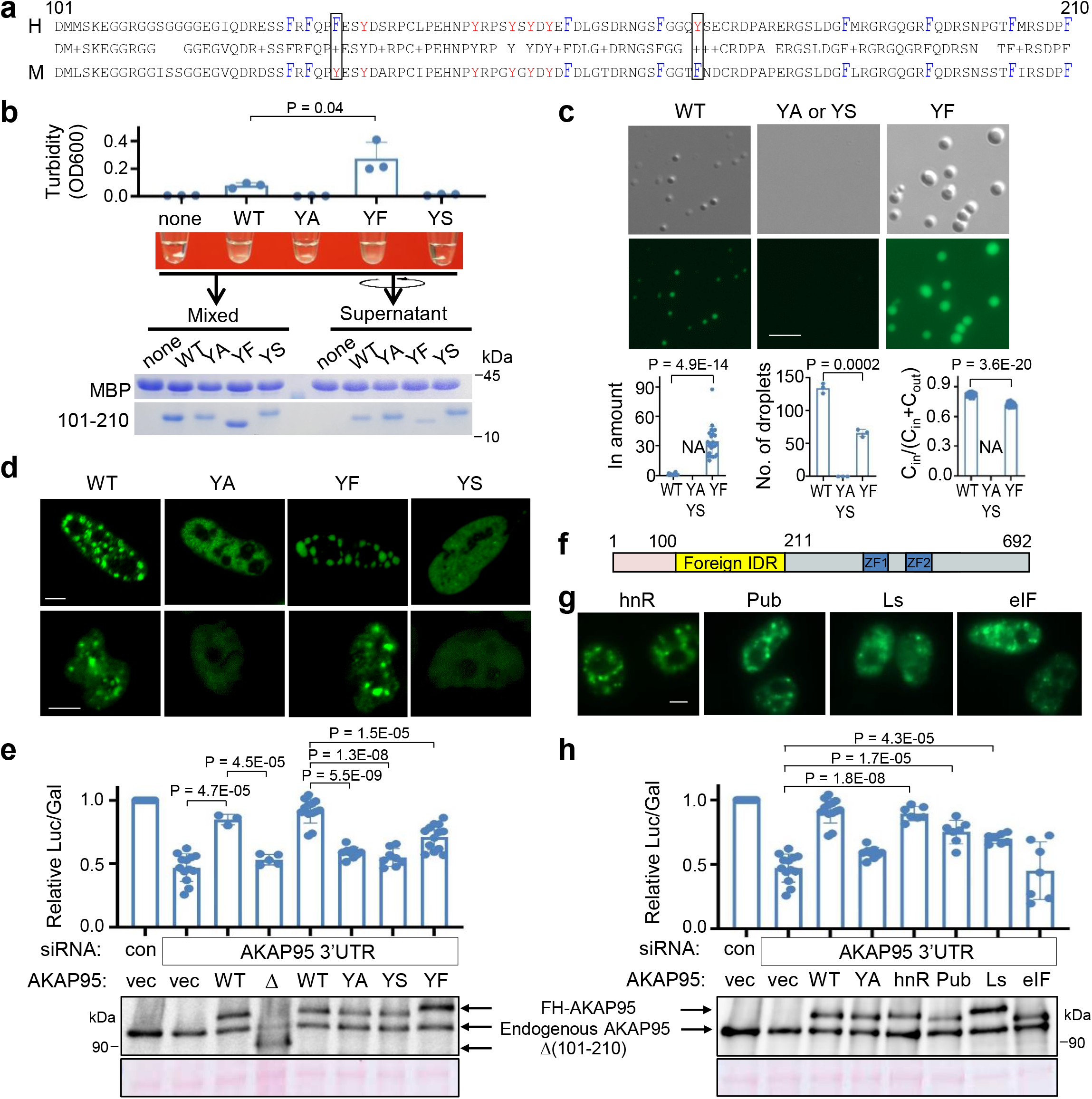
AKAP95 condensation requires Tyrosine in 101-210 and regulates splicing. **a,** Alignment of human and mouse AKAP95 (101-210). Middle row shows identical residues (by letter) and conservative mutations (by “+”). Tyr, red; Phe, blue and tall. Box, Tyr and Phe swapping. **b,** MBP alone (none) or MBP-AKAP95 (101-210) WT or mutants all at 50 μM and in 30 mM NaCl after TEV protease treatment for 30 min. Turbidity of each reaction was shown in pictures, and by OD600 as mean ± SD from n = 3 (for YA, YS) or 4 (the rest) independent assays. Samples taken after mixing (“mixed”) and from supernatant after centrifugation were resolved by SDS-PAGE followed by coomassie blue staining. **c,** DIC and fluorescence microscopy images for 10 μM Oregon-green-labeled MBP-AKAP95 (101-210) WT and mutants in 30 mM NaCl after TEV protease treatment for 30 min. Plots from left to right at bottom show the relative protein amount in droplet, number of droplets in a field, and the ratio of protein concentration inside droplets over sum of inside and outside droplets, respectively. Calculated as mean ± SD from n = 24 randomly picked droplets for WT or YF, except for number of droplets from n = 3 randomly picked fields. NA, not applicable. **d,** Fluorescence microscopy images of HeLa cells transfected with (top) and Flp-In T-Rex 293 cell lines expressing (bottom) GFP fusions with full-length AKAP95 WT or mutants. Repeated > 3 times. **e,h,** HEK293 cells co-transfected with indicated siRNAs and plasmids were subject to splice reporter assay (top) and immunoblotting with α-AKAP95 (bottom). Δ = Δ(101-210). Mean ± SD from n = 8 [except 5 for Δ(101-210) and 13 for YF and 2^nd^ WT] independent transfections are plotted in e and 7 independent transfections in h. **f,** Schematic of AKAP95 chimeras. **g,** Fluorescence microscopy images of 293T cells transfected with indicated AKAP95 chimeras fused to GFP. Repeated > 3 times. P values by two-sided Student’s *t*-test for b and one-way ANOVA followed by Tukey’s post hoc test for e and h. Scale bar, 5 μm for all. Uncropped blots are provided as source data.

In a splice reporter assay^33^, AKAP95 KD reduced minigene splicing, which was rescued by restored expression of WT, but not by mutants lacking 101-210 or containing YA or YS in 101-210 (**Fig. 6e**). Deletion of 101-210 did not affect binding of AKAP95 to DDX5 or hnRNP M (**Extended Data Fig. 6h**), key factors in splice regulation^33^. Moreover, an AKAP95 mutant lacking 1-386 is fully active in mediating chromatin condensation^45^, suggesting that Δ(101-210) does not grossly disrupt the protein structure. These results indicate that Tyr residues in 101-210 are crucial for AKAP95’s activity in splice regulation, likely through their critical role in mediating AKAP95 phase separation.

We reasoned that if 102-210 mediates splice regulation via its condensation property, we may be able to retain the splicing activity by replacing this region with a different condensation-capable region. We thus generated four chimeric proteins by replacing 101-210 of AKAP95 with IDRs that (i) have comparable sizes, (ii) are from proteins in pathways largely unrelated to AKAP95, and (iii) have been previously characterized to phase separate under proper conditions in vitro (**Fig. 6f, Supplementary Table 3**)^51^. hnRNP A1_IDR_ and eIF4GII_IDR_ form liquid-like condensates, while Lsm4_IDR_ tends to form fiber-like aggregates. Pub1_IDR_ forms condensates when coupled with an RNA-binding region and in the presence of RNA^51^. We found that all of these chimeras restored the punctate nuclear distribution when expressed at a level lower than endogenous AKAP95, and AKAP95-hnRNPA1_IDR_, −Pub1_IDR_, and −Lsm4_IDR_ chimeras all significantly rescued the reporter splicing (**Fig. 6h,g, Extended Data Fig. 6i**). These results strongly support the functional role of the phase separation property in AKAP95’s activity in splice regulation.

### The appropriate biophysical properties of AKAP95 condensates are important for splice regulation

As the YF mutation in 101-210 did not impair AKAP95 condensation, we were surprised that this mutant significantly impaired splicing (**Fig. 6e**). We suspected that YF may be more prone to form condensates than WT, as we observed higher turbidity, more portion in the condensates and the pellet fraction, and larger but less mobile droplets for 101-210 (YF) compared to 101-210 (WT) (**Fig. 6b,c, Extended Data Fig. 6d**). Indeed, YF reached much higher maximum turbidity and with faster rate than WT (**Extended Data Fig. 7a**). While condensation of both was promoted by increasing protein concentration and inhibited by increasing salt concentration, YF formed condensates at a lower protein and higher salt concentration than WT (**Fig. 7a, Extended Data Fig. 7b, c**). Consistent with the feature of phase separation, protein concentration in the dense phase and its ratio in light and dense phases remained largely constant as the total protein concentration increased above the saturation concentration (which was higher for WT than YF) (**Extended Data Fig. 7b, Supplementary Table 4**). While pre-formed YF condensates were also dissolved by high salt and re-formed upon dilution like WT, the re-formed YF condensates appeared more aggregated than WT (**Extended Data Fig. 7d**). Unlike the WT condensates that maintained a spherical morphology over time after MBP cleavage, YF appeared spherical at the early time point, but gradually developed into larger clusters with irregular morphology, suggesting a liquid-to-solid transition (**Fig. 7b**). Moreover, the condensates of WT, but not YF, could completely fuse into a spherical condensate (**Extended Data Fig. 7e, Supplementary Movies 4-6**).

FRAP assays showed YF recovered much less than WT within the assay time in vitro. Extended incubation allowed both to age and lose dynamics, with YF becoming entirely immobile (**Fig. 7c)**. Full-length YF also recovered much less than WT in nucleus (**Fig. 7d**). Moreover, Line Raster Image Correlation Spectroscopy (RICS) (**Extended Data Fig. 7f-k**) showed that YF significantly reduced the diffusion rate in the condensates in vitro (WT = 0.14+0.09 μm^2^/s, YF = 0.08+0.06 μm^2^/s) and in live cell nucleus (WT = 0.016+0.01 μm^2^/s, YF = 0.009+0.007 μm^2^/s) (**Fig. 7e-h**).

Taken together, these results strongly suggest that the YF mutation promotes a transition toward a more solid-like state with reduced dynamics and diffusion, which harms AKAP95’s activity in splice regulation.

### AKAP95 condensates with appropriate material properties regulate tumorigenesis

To study the role of AKAP95 condensation in tumorigenesis, we expressed full-length AKAP95 WT or mutants (YS or YF in 101-210) all at near endogenous levels in the AKAP95-KD MDA-MB-231 cells (**Fig. 8a**). Compared to the vector control, WT significantly enhanced the colony formation and liquid culture growth of AKAP95-KD cells. In contrast, YS had no effect. YF only modestly enhanced colony formation and growth, and the effects were significantly lower than WT (**Fig. 8b**,**c**). *SMAD6* expression was fully restored by re-introduction of WT, but not by YS or YF (**Fig. 8d**). A similar trend was seen on *CCNA2* expression (**Extended Data Fig. 8a**). Splicing of *CCNA2* intron 1 was significantly improved by WT, but not at all by YS, and only weakly by YF (**Fig. 8e**). Similar effects were seen on skipping of the alternative exons in *RPUSD3* (**Fig. 8f**) and *PPM1K* (**Extended Data Fig. 8b**).

When expressed at comparable levels in the MYC-transduced *Akap95* KO MEFs (**Fig. 8g**), AKAP95 WT, but not YS or YF, suppressed senescence (**Fig. 8h**). As shown by the analyses of global gene expression (**Extended Data Fig. 8c, Supplementary Table 2, tab 4**) and splicing (**Fig. 8j, Supplementary Table 2, tab 5**), compared to the vector control, YS minimally altered expression and splicing of genes that were affected by WT. YF substantially altered many genes but not as effectively as WT on a subset of genes. Many genes promoting cell proliferation and survival were upregulated by WT, but not YS or YF (**Fig. 8i**). Moreover, expression of certain SASP genes was suppressed by WT, but somewhat less effectively by YS or YF (**Extended Data Fig. 8d**). WT, but not YS, remedied the alternative exon inclusion on *Aamdc*, *Zfp518b, Numb*, and *Ttc21b* pre-mRNAs. While YF occasionally effected alternative splicing (e.g. for *Zfp518b*), it was overall much less effective than WT (**Fig. 8k, Extended Data Fig. 8e,f**). Taken together, these results indicate that the biological abilities of AKAP95 in promoting cancer cell growth and overcoming oncogene-induced senescence, and its biochemical activities in regulating gene expression and splicing, require its condensation into dynamic and liquid-like droplets, and is impaired by perturbation toward a more solid-like state (**Fig. 8l**).

## DISCUSSION

Our results allow us to propose a model for the role of AKAP95 phase separation in spatial regulation of gene expression that underlies tumorigenesis (**Extended Data Fig. 8g**). AKAP95 binds to many transcription and splicing regulators at its N-terminus, and to RNAs at its ZFs. Its 101-210 drives condensation, thereby creating a micro-compartment with factors at high local concentration and appropriate material properties to ensure optimal activities for gene expression. Our data link the biophysical properties of AKAP95 condensates to its physiological function in tumorigenesis via its biochemical activities in gene regulation.

Cancer cells hijack normal pathways to cope with numerous stresses not encountered by normal cells and become exceptionally dependent on these pathways^13, 52^. MYC selectively amplifies expression of numerous genes^53^ and greatly enhances the burden on many pathways including transcription and splicing, thereby sensitizing cells to even partial inactivation of these pathways^54–56^. Indeed, *MYC* greatly promotes the expression of *Akap95* but not its more ancient paralog, *Akap8l*, and requires elevated AKAP95 for efficient expression of many MYC targets (**Figs. 3h,i, Extended Data Fig. 3f,h**). We surmise that the newly evolved AKAP95 joins AKAP8L in regulating gene expression for cell growth and survival, ensuring genetic robustness in normal cells. The elevated demand for gene expression and tumor-suppressive responses in cancer cells call for elevated levels of AKAP95 and its related factors, and the preferential upregulation of AKAP95 by the oncogene makes cancer cells particularly vulnerable to AKAP95 depletion. AKAP95 is thus a potential target for cancer treatment with a therapeutic window.

Our results indicate that AKAP95 condensation requires the aromatic ring of Tyr, with probable contributions from cation-π and π-π interactions^57^, but is weakened by the Tyr hydroxy group. It is thus important to maintain a balance of Phe and Tyr in a protein for proper material state and biological activities, as partly reflected by the conservation and mutual swap of these two residues in AKAP95 (101-210) between mouse and human (**Fig. 6a**). The opposite effects of YF mutation on AKAP95 to that on the FUS family proteins^50^ suggests a context-dependent regulation of condensate properties by the same amino acids.

We have provided several lines of evidence, including mutations and IDR-chimeras that disrupt and regain condensation properties, to support a causal role of AKAP95 phase separation in its biological activities. We note that, while its IDR completely rescues splicing activity, hnRNP A1 is neither physically nor functionally (in splicing regulation) associated with AKAP95^33^. The low splicing abilities of other IDR chimeras can be due to multiple reasons. Lsm4_IDR_-chimera showed modest splicing activity comparable to AKAP95 (YF), consistent with both Lsm4_IDR51_ and AKAP95 (YF) prone to form less dynamic condensates. The inconsistent splicing activity of the eIF4GII_IDR_-chimera is likely because the yeast eIF4GII_IDR_ has intrinsic RNA-binding activity^51^, which would conceivably interfere with binding of the chimera to correct RNA regions required for splice regulation in human cells.

Complete inactivation of AKAP95 by YS or YA mutation probably results from failure in concentrating key factors into local proximity. On the other hand, the hardened YF condensates may slow down the biochemical reaction kinetics inside by restricting the movement and interactions of other macromolecules such as other splicing modulators and RNA substrates. The often imperfect spherical shape and incomplete recovery after photobleaching suggest that WT AKAP95 is probably somewhere between a highly viscous liquid and a hydrogel state (**Fig. 8l**). Liquidity of high-order assemblies requires dynamic binding modes arising from multivalent interactions with low affinity and flexible connecting regions that is often disordered and contributes to intrinsic fuzziness^58, 59^. Our data suggest that tethering to RNAs may restrict AKAP95 dynamics, but it remains unclear how disordered and flexible AKAP95 is when engaged with other macromolecules in the cell and how this affects its activity. As exemplified by nucleolus, the most prominent non-membrane organelle in cell^60–62^, further studies will help better understand how the quantitative variations in the biophysical properties of the AKAP95 condensates produce different outcomes in gene expression and disease.

The partial co-localization of AKAP95 condensates with actively transcribing Pol II in nucleus supports their direct role in mRNA biogenesis, likely including transcription and splicing. The dynamic liquid droplets may also provide a proper microenvironment for assembly, modifications, and dynamic supply of factors needed for active splicing at other locations, consistent with the partial co-localization of AKAP95 with nuclear speckles known to have these functions^47^. Our findings suggest that, in addition to the two simple states (diffuse or condensate) that act as an on/off switch for activity^18–22^, changes in dynamics and fluidity of a gene regulator in the continuum of material property can significantly impact its biological activity (**Fig. 8l**). Our findings also raise the unconventional possibility that cancer may be inhibited by perturbing the material properties of the condensates of key players in either direction, i.e., disrupting or hardening the condensates (**Fig. 8l**).

## Methods

### Animals

All animal procedures were approved by the Institutional Animal Care and Use Committee at the University of Alabama at Birmingham. All mice were maintained under specific pathogen–free conditions and housed in individually ventilated cages. Immune-deficient NSG (NOD.Cg-Prkdcscid Il2rgtm1Wjl/SzJ) mice were purchased from The Jackson Laboratory. To generate the *Akap95* KO mice, embryonic stem cells on C57BL/6 background that harbor an Akap8^tm1a(EUCOMM)Hmgu^ (Knockout-first, thus null “-“) allele were purchased from EUCOMM (Helmholtz Zentrum München, Germany), and were injected into blastocysts to generate chimeras at the University of Alabama at Birmingham Transgenic Mouse Facility. *Akap95^+/−^* mice with germline transmitted *Akap95*-targeted allele (−) allele were selected for further crossing to generate WT, *Akap95^+/−^ and Akap95^−/−^* mice. Peripheral blood profiles were measured using Hemavet 950 (DREW Scientific Inc)

### Cells

Primary mouse embryonic fibroblasts (MEFs) isolation, culture, growth assays, transformation, soft agar colony formation, and in vivo tumorigenesis assays were performed as previously described^54^. Briefly, MEFs were harvested from E13.5 embryos from cross of *Akap95*^+/−^ female and male mice. Early passages of MEFs were frozen in the liquid nitrogen, and cultured in DMEM plus 10% FBS at use. Flp-In T-REx 293 cells stably expressing AKAP95 WT or mutants were generated using the Flp-In system (Thermo Fisher Scientific) according to the manufacturer’s instructions. Protein expression was induced with doxycycline for 48 hrs. MDA-MB-231 cells were a kind gift from Lizhong Wang (University of Alabama at Birmingham). HeLa, MDA-MB-231, MCF7, HEK293, 239T, Flp-In T-REx 293 cells were cultured in DMEM (Thermo Fisher Scientific) supplemented with 10% fetal bovine serum (Thermo Fisher Scientific).

### Constructs

Human *AKAP95* cDNA in pcDNA5/FRT/TO (Invitrogen) with an N-terminal FlAG-HA-(FH-) tag was previously described^32^. Some constructs have EGFP (all called GFP in this study) fused at the C-terminus. Truncation mutants of AKAP95 were generated by PCR cloning. In particular, the FH-AKAP95 Δ(101-210) construct was generated with a BamHI site replacing the 101-210 region. Tyrosine mutants of the 101-210 region in AKAP95 were synthesized (Biomatik USA, LLC, Delaware, USA). The full length AKAP95 mutants were achieved through in-fusion PCR cloning. To generate AKAP95 chimeras, human hnRNPA1 (186-320), yeast Pub1 (243-327), yeast Lsm4 (91-187), and yeast eIF4GII (13-97) sequences were cloned from the cDNAs of human HEK293 cells and yeast Saccharomyces cerevisiae, respectively, and inserted at the BamHI site in the FH-AKAP95 Δ(101-210) construct to generate the AKAP95 chimeras. For rescue constructs, AKAP95 WT and full-length AKAP95 mutants that have indicated tyrosine mutations within 101-210 were cloned into pTripZ (Dharmacon) between AgeI and MluI sites (replacing the tRFP and mir sequences in the vector) for doxycycline-inducible expression, and also cloned into pCDH-MSCV-MCS-EF1α-GFP+Puro lentiviral vector (System Biosciences, Cat# CD713B-1, Palo Alto, CA) for constitutive expression. For protein expression as MBP-fusion, AKAP95 or its variants and FUS, sometimes with GFP fused to the N-terminus of AKAP95 regions (as MBP-TEV site-GFP-AKAP region) as indicated in the figure legends, were cloned into pET28-MBP-TEV vector (Addgene: #69929) downstream of the TEV cleavage site. The sequences of all plasmids were confirmed by Sanger sequencing.

### Antibodies

Rabbit polyclonal anti-AKAP95, Santa Cruz Biotechnology Cat#sc-10766; Rabbit polyclonal anti-AKAP95, Bethyl Laboratories Cat#A301-062A, Mouse monoclonal anti-GAPDH, EMD Millipore Cat#MAB374; AlexaFluor 555 conjugated goat anti-rabbit IgG, Thermo Fisher Scientific Cat#A-21428; Mouse anti-SRSF2 antibody, Abcam Cat#Ab11826; mouse anti-Pol II, 8WG16 clone, COVANCE Cat# MPY-127R; Mouse anti-pol II CTD-S2P, H5 clone, COVANCE Cat#MPYT-127R; Rabbit polyclonal anti-cyclin A, Santa Cruz Biotechnology Cat#sc-751; Rabbit polyclonal anti-DDX5, Santa Cruz Biotechnology Cat#sc-32858; Mouse monoclonal anti-hnRNP M, Santa Cruz Biotechnology Cat#sc-20002; AlexaFluor 555 conjugated goat anti-mouse IgG, Thermo Fisher Scientific Cat# A-21422. A dilution of 1:1000 was used for immunoblotting except 1:2000 for GAPDH.

### Virus package and concentration

Lentiviral vectors were packed with psPAX2 and pMD2.G in 293T cells with polyethylenimine (PEI) transfection reagent. Retroviral vectors pBabe-puro Ras V12 (Addgene #1768) and pWZL-Blast-myc (Addgene #10674) were packed with pEco (Clontech) in 293T cells. The viral solutions were 30-fold concentrated by PEG6000 precipitation, then aliquot and stored at −80°C.

### Gene knockdown, cell growth, colony formation, and assays to determine transcript splicing and expression levels

AKAP95 KD was performed by either lentivirus-mediated shRNA or transient transfection of siRNA as described ^33^. Detailed information for shRNAs and siRNAs is in Data S3. MDA-MB-231 cells infected with pLKO-based shRNA lentivirus were selected under 20 μg/ml blasticidin for 3 days. For rescue assays, cells were infected with pCDH-based lentivirus and selected in 2 μg/ml puromycin for 3 days, and further infected with pLKO-based shRNA lentiviruses for KD. For cell growth assays, 5 × 10^4^ cells/well seeded in 6-well plates were cultured in DMEM plus 10% FBS without antibiotics, cell number was counted manually at 5 days of culture or at indicated time points for multiple-time points assays, and cells were fixed with 4% formaldehyde and stained with 0.1% crystal violet solution 5 days after culture. For colony formation assay, 3,000 cells/well in 6-well plates were cultured without antibiotics, and cells were fixed with 4% formaldehyde and stained with 0.1% crystal violet solution and quantified after10-day culture.

MDA-MB-231 scramble or *AKAP95* knockdown cells were treated by 50 μg/ml of cycloheximide (Millipore Sigma) for 6 hours or transfected by siUPF1 or siBTZ to inhibit NMD. Total RNA were harvested for reverse transcription with random hexamer primers. *CCNA2* transcript isoforms were detected by regular PCR with the splicing primers listed in Supplementary Table 3. To characterize the stability of *CCNA2* in cells, MDA-MB-231 scramble or *AKAP95* knockdown cells were treated with 5 μg/ml of Actinomycin D (Millipore Sigma) or Actinomycin D together with cycloheximide for 6 hours. At indicated time points, total RNA were harvested and *CCNA2* transcript relative amounts were measured by qPCR with primers listed in Supplementary Table 3.

### Cell proliferation assay and apoptosis assay

Cell proliferation (by BrdU incorporation) and apoptosis (by Annexin V staining) assays were performed as previously described^63^. For cell proliferation assay, control and AKAP95-KD MDA-MB-231 cells were incubated with BrdU at a final concentration of 10 μM in cell culture medium for 45 min. Cells were harvested and then processed with BrdU staining using the FITC-BrdU Flow kit (BD Pharmingen) following the manufacturer’s instructions. For apoptosis assay, cells were harvested and washed twice with cold PBS containing 3% heat-inactivated FBS. Cells were then incubated with FITC-Annexin V (BioLegend) and 7-amino-actinomycin D (7-AAD) for 15 minutes in binding buffer (10 mM HEPES, 140 mM NaCl and 2.5mM CaCl_2_) at room temperature in dark. The stained cells were analyzed immediately by flow cytometry on LSRFortessa (Becton Dickinson). Data were analyzed using the FlowJo software.

### MEF transformation assay and senescence assay

MEFs in early passage (less than four passages) were infected with *H-RAS^G12V^* and c-MYC viral particles followed by selection in puromycin (2 μg/ml) and blasticidin (10 μg/mL). Soft agar colony formation assays were performed by plating transformed MEFs in a 6-well plate at 2,500 cells/well. MEFs were cultured in a layer of MEF culture medium in 0.35% agar over a base layer composed of culture media in 0.5% agar and fed every 4 days. Colonies were formed over the course of 3 to 4 weeks. For MYC-induced senescence assay, MEFs were infected with c-MYC retrovirus and selected against 10 μg/mL blasticidin for 3 days. Four to 5 days post selection, 10^5^ cells/well were seeded in 6-well plate. Twenty-four hrs later, the cells were washed twice with PBS, fixed and stained for β-galactosidase activity at pH = 6 using Senescence Cells Histochemical Staining Kit (Sigma, Cat#CS0030). For rescue assays, MYC-transduced *Akap95^−/−^* MEF were infected with AKAP95- or mutant-expressing pTripZ lentiviruses and selected under 2 μg/ml puromycin for 3 days. After selection, cells were cultured under 100 ng/mL doxycycline for 4-5 days and then stained for β-galactosidase activity.

### Allogeneic and xenogeneic transplantation

Transformed MEF cells and MDA-MB-231 cells were washed and re-suspended in PBS. Subcutaneous injection of 2 × 10^6^ MDA-MB-231 cells in 100 μl PBS were administered into left and right flanks of 8-week-old female NSG mice (NOD.Cg-*Prkdc^scid^ Il2rg^tm1Wjl^*/SzJ, Jackson Laboratory, stock No. 005557), respectively. The same amount of transformed MEF were inoculated into left and right flanks of 8-week-old male NSG mice in the same way. Tumor size was measured in the 2 longest dimensions using a Vernier caliper. Tumor volume (V) was calculated with the formula V = D1(D2)^2^/2, where D1 is the long dimension and D2 is the short dimension. Four weeks after transplantation, mice were humanely sacrificed for collecting tumors. The tumors were also weighed.

### Analyses for protein disorder, domain, and amino acid composition

Disordered regions were identified using IUPred (http://iupred.elte.hu/). Amino acid composition was analyzed by Composition Profiler (http://www.cprofiler.org/cgi-bin/profiler.cgi) by using SwissProt 51 Dataset as the background sample.

### Co-immunoprecipitation assay

Co-immunoprecipitation assays were performed as described^33^. Briefly, 293T cells were transfected with indicated constructs. Twenty-four hrs later, cells were lysed in BC300 [50mM Tris (pH 7.4), 300mM KCl, 20% glycerol, 0.2 mM EDTA] with 0.1% NP40, 1mM DTT, and protease inhibitor cocktail (Roche, Cat# 4693159001). Lysates were incubated with the anti-Flag M2 antibody (Sigma, A2220) and washed by the lysis buffer. Bead-bound proteins were resolved by SDS-PAGE and detected by immunoblotting using indicated antibodies.

### Protein expression and purification

The pET28-MBP-TEV-based constructs were transformed into BL21 Star (DE3) (Thermo Fisher Scientific, Cat# C601003) *E. coli*. Bacteria culture at OD600 of 0.6 were induced with 0.4 mM of Isopropyl β-D-1-thiogalactopyranoside (IPTG) for 5 hours at 25°C to express MBP-fusion proteins. The pET28-MBP-TEV vector itself was used to express MBP control, which was MBP and a small peptide (sequence GSLSTGCY) after cleavage by TEV protease. Bacterial cells were resuspended in BC500 [50 mM Tris (pH 7.4), 500 mM KCl, 20% glycerol, 0.2 mM EDTA], 0.1% NP40, protease inhibitor cocktail (Roche, Cat# 4693159001), 1 mM DTT, and lysed by sonication. After centrifugation, the supernatant was incubated with pre-equilibrated Amylose resin (New England BioLabs, Cat# E8021) in batch for three times. The resin was pooled, washed with the lysis buffer 4 times with one additional wash in BC0, and eluted in multiple fractions in 10 mM Maltose in BC0 buffer. Purified proteins were examined by SDS-PAGE followed by coomassie blue staining. Protein concentration was determined by Nanodrop measurement for OD_280_ and calculation using extinction coefficient provided by ExPASy ProtParam (https://web.expasy.org/protparam/), and also validated by comparing with coomassie blue staining of known concentration of BSA as well.

### Estimation of nuclear concentration of AKAP95

By immunoblotting of lysates from known number of cells alone with purified and quantified AKAP95, and based on nuclear volume as 220 fl^64^, we calculated the nuclear concentration of endogenous AKAP95 to be ~2.7 μM in 293 cells and ~2.1 μM in MDA-MB-231 cells.

### In vitro phase separation assay

For in vitro phase separation assays, the MBP-tag was cleaved off the fusion proteins by ProTEV Plus (Promega, Cat# V6102). Cleavage efficiency was typically ~70-90% at 30 min at room temperature, and showed modest increase as digestion time increased. Sometimes, AKAP95 (101-210) and its mutants were fluorescently labeled with Oregon green 488 carboxylic acid succinimidyl ester following manufacturer’s instructions. Briefly, the protein solution was first replaced with 0.1 M sodium bicarbonate buffer by dialysis. 20 μl of 10 mg/ml fluorophore were added into 2 mg protein and incubated at 4 °C in dark overnight. The protein was purified on a Bio-Spin® P-6 Gel Columns (BioRad) desalting column equilibrated in BC-0 buffer to separate labeled protein from excess fluorophore and then stored at −80 °C. The calculated labeling efficiency was 1 in 2 molecules for all proteins.

Phase separation was assessed by 4 ways: (1) visual observation of the turbidity. (2) Measurement of the optical density at wavelength of 600 nm (OD600) or 450 nm (OD450) for turbidity by Nanodrop. The kinetics of turbidity development was determined by OD450 measurement in 96-well plate on a Synergy HTX plate reader (BioTek). Readings were taken once every 10 seconds and analyzed by the Gen5 Data Analysis Software (BioTek). (3) Separation of the condensates from the aqueous phase by centrifugation at 5000 rpm for 10 min, followed by SDS-PAGE and Coomassie blue staining of the supernatant and mixed samples. (4) Microscopy-based method. After TEV protease treatment of indicated time, the protein mixture was added into a homemade flow chamber comprised of a glass slide sandwiched by a coverslip with one layer of double-sided tape as a spacer. Images were taken on Zeiss Axio Observer Z1 microscope or Zeiss LSM780 confocal microscope with 60X oil lens. Fluorescence intensity inside (*I*_in_) and outside (*I*_out_) of droplets as well as droplet diameters (*D*) were analyzed by ZEN2.6 software (Carl Zeiss). Assuming fluorescence intensity reflecting protein concentration, concentration ratio *C*_in_/(*C*_in_ + *C*_out_) is calculated as *I*_in_/(*I*_in_ + *I*_out_). Protein amount in each droplet were calculated as *I*_in_ ×4/3π(D/2)^3^.

### Immunofluorescence and live cell imaging

HeLa, MDA-MB-231, and MEF cells were washed with PBS and fixed with ice cold methanol for 15 min. The fixed cells were incubated with 1:250 rabbit anti-AKAP95 antibodies overnight at 4°C, and developed with 1: 300 AlexaFluor 555 conjugated goat anti-rabbit secondary antibody for 1 hr at room temperature. For AKAP95 colocalization with SRSF2, HeLa cells were transfected with pcDNA5-flag-HA-AKAP95-GFP 24 hrs before fixation. Cells were then incubated with mouse anti-SRSF2 (SC-35) antibodies followed by AlexaFluor 555 conjugated goat anti-mouse secondary antibody. The slides were further counterstained with DAPI. All immunostaining images were acquired on Nikon A1 confocal microscope with 60X oil lens. Intensities of AKAP95 and SRSF2 signals were analyzed by ImageJ. For live cell imaging, HeLa cells transiently transfected with indicated constructs and Flp-In T-REx 293 stable cells were cultured in 35 mm glass-bottom dishes (MatTek), and used for imaging on Zeiss LSM780 confocal microscope supported with a Chamlide TC temperature, humidity and CO_2_ chamber. Images were collected by either 40X or 60X oil lens.

### Fluorescence recovery after photo-bleaching (FRAP) assays

FRAP assays were performed on Zeiss LSM780 confocal microscope. Live cell FRAP assays were performed at 37°C, and the fluorescence signal was bleached using 40% of maximum laser power of a 488-nm laser for approximately 8 sec. For FRAP in vitro, Oregon-green 488 labeled proteins were spiked into unlabeled proteins, or GFP-tagged proteins were used. Samples were transferred to homemade chambered slides. In vitro condensates were bleached at the center with 100% of maximum laser power of a 488-nm laser for approximately 5 sec. For both in vitro and in vivo FRAP, bleaching area was 1-2 μm in diameter. The recovery was recorded at the rate of 2 s/frame. Condensates of similar size between samples were used in the same experiments. After subtraction of background signal, intensities were normalized for global photobleaching (from a neighboring unbleached droplet) during image acquisition according to^65^. FRAP recovery curves were fitted to generate recovery time for each bleached condensate, which was used to calculate mean and SD of the recovery time. Relative recovery was calculated as [I_max_-I_min_]/[I_0_-I_min_] x 100%, in which I_max_, I_min_, and I_0_ are the normalized maximum, minimum (0 sec after bleaching), and initial (before bleaching) intensities, respectively.

### Line RICS

To prepare for live cell Line RICS experiments, a 35 mm dish (MatTek, Ashland, MA, USA) was coated with Fibronectin (final concentration of 4 μg/ml in DPBS) for 2 h at 37°C and then plated with ~10^6^ HeLa cells. After 8 hr, the cells were transfected using the Lipofectamine 2000 kit (2 μl) and OPTI-MEM (100 μl) with 200 ng of plasmid, and mixed in 1 ml of DMEM. The cells were kept in the 37°C and 5% CO_2_ incubator overnight and then placed on the microscope which was equipped with an incubation system set at 37°C, 5% CO_2_.

To prepare for in vitro Line RICS experiments, 7.4 μl of 13.5 μM MBP-fused protein was mixed with 0.5 μl of 20X TEV buffer, 0.1 μl of 100 mM DTT, 1 μl TEV Protease, and 1 μl of 1.2 M NaCl. The solution was kept for 30 min at 25°C to allow the TEV protease to cut the MBP tag. After that 8 μl of the sample was taken to mix with 2 μl of 50% PEG6000. After 5 min, 2 μl of this mixture was placed onto home-made slide chambers (diameter 4 mm, thickness 0.1 mm plastic) in the between a glass slide and a coverslip glass. Finally, the chamber was placed on the sample holder for imaging.

All Line RICS autocorrelation measurements were performed using a Zeiss LSM 880 microscope equipped with Quasar where the emission signal is spectrally separated by passing through a grating onto the photomultiplier tubes (PMTs). An argon laser (Melles-Griot, Carlsbad, CA, USA) was set at 488nm excitation with a 488 primary dichroic mirror and a Zeiss Plan Apochromat 63x, 1.4 NA DIC M27, oil-immersion objective. The Line RICS acquisition has been carried out sampling a linear region across the condensates of 6 μm with ~10^4^ lines at 128 pixels in the x-axis, with a pixel size of 50 nm and scan line time of 0.101 ms/line. The temporal line carpet collected as a function of time was divided by ~64 frames of 128 pixels x 128 lines and analyzed with Global for Images SimFCS software (Version 4, Laboratory for Fluorescence Dynamics, Irvine, CA, USA). We used the line RICS algorithm with a moving average of 10 frames to correct the photobleaching and particle movement as reported previously^66^. The Line RICS algorithm computes the autocorrelation line by line, averaging the autocorrelation for each frame and then for the entire data set, producing a final autocorrelation function. Frames with large particles motion were discarded for the computation of the autocorrelation curve. The line RICS autocorrelation curves were fitted with a 1-specie Brownian diffusion fitting model as reported in the Supplementary Information.

### Splicing reporter assay

The double reporter splicing assay was performed on pTN24 reporter plasmid^67^ as described^33^. Briefly, HEK293 cells were transfected with relevant plasmids or siRNA using Lipofectamine 2000 reagent (Thermo Fisher Scientific), and harvested 48 h after transfection. Beta-galactosidase and luciferase activities were measured using Dual-Light System (Applied Biosystems).

### Cancer database analysis

AKAP95 association with human cancer was analyzed by cBioPortal (http://www.cbioportal.org/). RNA-Seq–derived gene expression levels from The Cancer Genome Atlas (TCGA) were analyzed by UALCAN portal^68^ at http://ualcan.path.uab.edu. Out of the 31 cancer types at the portal, 28 cancer types were analyzed as they can be clearly sub-grouped based on cancer stages, tumor grade.

### RT-PCR and RT-qPCR, and genome-wide expression and splicing analyses

RT-PCR and RT-qPCR to examine relative mRNA levels and alternative splicing, respectively, were performed as previously described^33^. For RT-PCR, briefly, alternative splicing on selected targets was determined by PCR, and mRNA levels were determined by qPCR, both following RNA isolation and reverse-transcription. Primers are listed in Data S3.

Total RNA quality was assessed with an Agilent 2100 Bioanalyzer. Samples with RNA integrity number greater than 9 were further processed to library preparation and sequencing. For RNA-seq, 2 rounds of polyA+ selection were performed and followed by conversion of mRNA to cDNA. Illumina sequencing adapters and barcodes were then added to the cDNA via PCR amplification. Indexed libraries were then normalized and pooled. 2 x 150 pair-end sequencing was performed using the HiSeq2500 (Illumina) with recommended protocol. 40-60 million reads per sample were mapped to human or mouse reference genome (hg19 and mm9, respectively) using Tophat version v2.0.4. Transcript assembly was performed using HTseq v0.10.0. Assembled RNA-seq count files were imported into RStudio software for analysis of differential expression with DESeq2 v3.8 package. Differential expressed genes with mean count across all groups greater than 10 were chosen for visualization with values scaled by row using RStudio. Gene ontology analysis and Pathway analysis was conducted with DAVID v6.8 and GSEA (Broad Institute, Cambridge, MA, USA) software packages. All GSEA was carried out by using the Hallmark gene sets from the Molecular Signature Database (MsigDB) except that the “Senescence_up” gene set^69^, a collection of genes upregulated in cells undergoing senescence, and the SASP gene set^44^ were curated from literature.

Alternative splicing isoforms were analyzed with MISO software^70^ with its version 2.0 annotation of all known AS events according to the online documentation (https://miso.readthedocs.io/en/fastmiso/). Briefly, accepted alignments from Tophat2 output within each sample were imported to MISO software with command miso –run with paired-end option included. Next, pairwise comparisons between samples were performed using compare_miso - compare-samples to detect differentially expressed isoforms. Five different type of alternative splicing events were analyzed: skipped exons (SE), alternative 3’/5’splice sites (A3SS, A5SS), mutually exclusive exons (MXE) and retained introns (RI). As SE was found to be the major type in all analyses, we chose to focus on SE for our detailed analyses in splicing heatmaps. MISO outputs were filtered according to the percent-spliced-in (PSI) values to quantify isoform expression.

### Quantification and statistical analysis

Statistical parameters including the definitions and exact values of n (e.g., number of experiments, number of cells, number of colonies, etc), distributions and deviations are reported in the Figures and corresponding Figure Legends. All data were expressed as mean ± SD; n.s., not significant, *P < 0.05, **P< 0.01 and ***P < 0.001 using one-way ANOVA with Tukey’s post hoc test (for multi-sample groups), 2-sided unpaired Student’s *t*-test or Mann-Whitney *U* test (for two-sample comparison and certain pair-wise comparisons to a specific sample in a multiple sample group), or log-rank test (for survival analysis), as indicated in Figure Legends. Statistical analysis was performed by in SPSS (IBM), Prism-GraphPad, or Excel. A P value of less than 0.05 was considered significant.

## Data availability

The RNA-seq data have been deposited in Gene Expression Omnibus database with the accession number GSE122308.

## Acknowledgments

We thank Lizhong Wang for providing the MDA-MB-231 cells, and Ying Gai Tusing for technical assistance with mice work. This work was supported by Start-up funds from the University of Alabama at Birmingham and the University of Virginia, and a Department of Defense Breast Cancer Research Program Breakthrough Award (BC190343). The confocal microscopy system at the Keck Center of University of Virginia was supported by a grant from NIH (OD016446; PI: Ammasi Periasamy). H.J. is a recipient of the American Society of Hematology Scholar Award, the American Cancer Society Research Scholar Award (RSG-15-166-01-DMC), and the Leukemia & Lymphoma Society Scholar Award (1354-19). F.P. and M.A.D. were supported in part by a grant from the NSF (MCB-1615701). M.A.D. and E.G. were funded by NIH grant number P41-GM103540.

## Author Information

These authors contributed equally: Wei Li, Jing Hu, Bi Shi.

## Contributions

H.J. conceived, designed, and supervised the study, and wrote the manuscript. W.L. designed and performed molecular and functional analyses for tumorigenesis, and most studies on protein condensation in vitro and in vivo, and also participated manuscript writing and editing. J. H. made most constructs and cell lines, initiated and participated the protein condensation assays in vitro and in vivo, and performed molecular and functional analyses. B.S. conducted all bioinformatic analyses and performed molecular analyses. F.P. performed and analyzed the RICS experiments under the guidance of M.A.D. and E.G.

## Competing interests

The authors declare no competing interests.

## Supplementary Information

### Extended Data Figure Legends

**Extended Data Figure 1.**
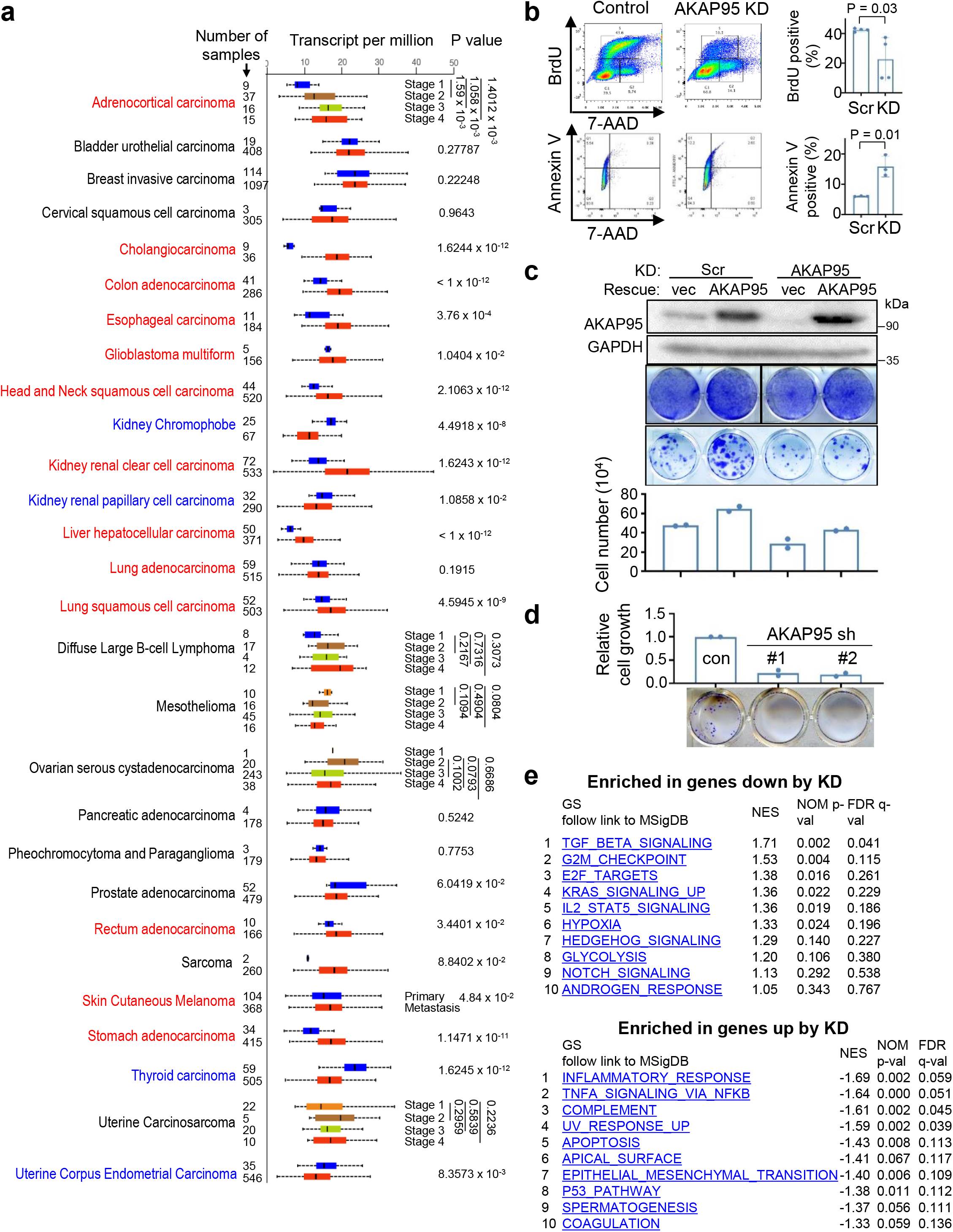
AKAP95 regulates cancer cell growth and gene expression. **a,** RNA-Seq–derived gene expression levels from TCGA were analyzed by UALCAN portal. Box plot analysis shows relative expression of *AKAP95* in 28 types of cancer (red box) versus normal (blue box) samples, unless indicated for different cancer stages or tumor grades for certain cancer types. Cancer type in red font has significantly higher AKAP95 expression in cancer than in normal (or in later stage than in earlier stage). Cancer type in blue font has significantly lower AKAP95 expression in cancer than in normal samples (or in later stage than in earlier stage). Numbers on the left of the plot stand for the number of samples. Data are median (line), 25–75th percentiles (box) and minimum-maximum values recorded (whiskers). **b,** Assays for cell proliferation by BrdU incorporation (top) and apoptosis by Annexin V staining (bottom) for control and AKAP95-KD MDA-MB-231 cells. Images of flow cytometry results are shown (left), and percentages of cells positive for BrdU or Annexin V are shown as mean ± SD from n = 3 independent KD assays. **c,** MDA-MB-231 cells were infected to express scramble (control) or AKAP95 shRNA #1 (KD) and indicated constructs. Top, immunoblotting of total cell lysates. Middle, images of these cells seeded at high (5 ×10^4^ cells/well in 6-well plate, top) and low (400 cells/well in 24-well plate, bottom) densities and stained with crystal violet. Bottom, cell numbers (seeded at high densities) as mean ± SD from n = 2 independent experiments. **d,** MCF7 cells were infected to express scramble control shRNA or two AKAP95 shRNAs. Bottom, images of indicated cell colonies stained with crystal violet. Top, cell numbers were quantified and presented in the bar graph as mean ± SD from n = 3 independent experiments. **e,** Top 10 gene sets enriched in genes down-(top, n = 951) and up-(bottom, n = 294) regulated by AKAP95 KD in MDA-MB-231 cells. NES, normalized enrichment score. P values by two-sided Student’s *t*-test for a-d and modified Fisher’s exact test for e. Uncropped blots are provided as source data.

**Extended Data Figure 2.**
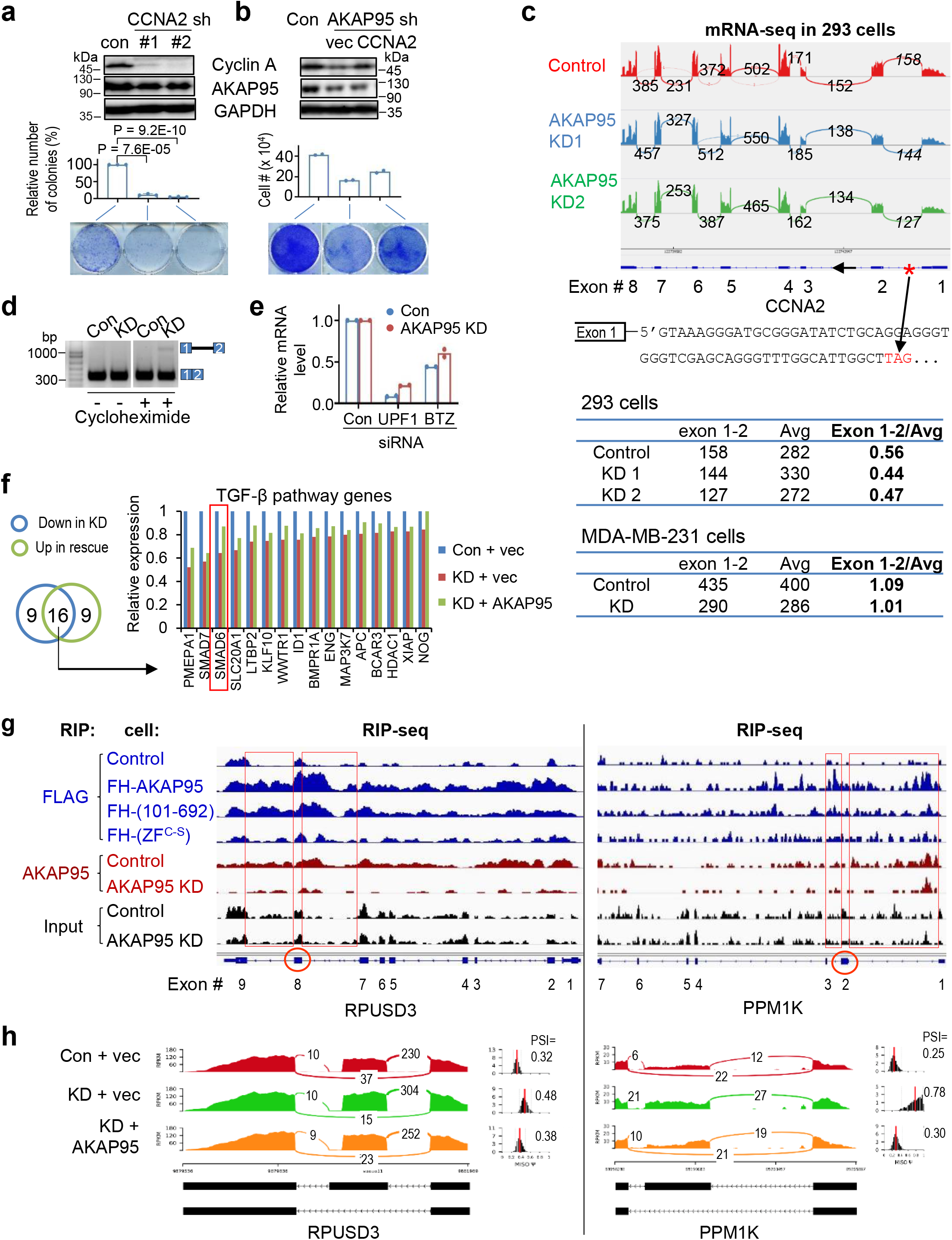
AKAP95 directly regulates splicing of key transcripts for cancer. **a,b,** MDA-MB-231 cells virally expressing control or indicated shRNAs (**a**) or shRNA combined with indicated constructs (**b**) were subject to immunoblotting of total cell lysates (top) and colony formation assay. Middle, colony numbers as mean ± SD from n = 3 (**a**) or 2 (**b**) independent experiments. Bottom, images of cells stained with crystal violet. **c,** Top, mRNA-seq profiles for *CCNA2* in control or 293 cells of AKAP95 KD^1^. The numbers of exon junction reads are indicated. The red asterisk at the gene diagram indicates a stop codon 57 bp downstream of exon 1 in the intron. The number of reads for the junction of exons 1 and 2, and for the average neighboring exons, and their ratios are in the tables below for indicated cells. **d,** Total RNAs were used for RT-PCR for intron 1 region in control and AKAP95-KD MDA-MB-231 cells treated with or without cycloheximide for 6 hours. Repeated > 3 times. **e,** Relative mRNA levels of *UPF* in UPF1-KD samples and *BTZ* in BTZ-KD samples, respectively, each relative to the control samples, as determined by RT-qPCR and normalized to *GAPDH*. **f,** Relative expression level of TGF-β pathway genes based on RNA-seq reads from control and AKAP95-KD MDA-MB-231 cells expressing vector or AKAP95. Venn diagram shows numbers of TGF-β pathway genes (from GSEA) downregulated by AKAP95 KD and upregulated by rescue with AKAP95 expression, and the relative expression of the 16 overlapped genes in both categories are plotted. **g,** RIP-seq profiles showing AKAP95 binding to *RPUSD3* and *PPM1K* pre-mRNAs. Track information is the same as in Fig. 2c. Red circles indicate the alternatively included exons (corresponding to the middle exon in the gene diagrams in (**h**), and red boxes show AKAP95 binding at the introns flanking these exons. **h,** Sashimi plots showing that the alternative splicing of *RPUSD3* and *PPM1K* pre-mRNAs was affected by AKAP95 KD and rescued by restored expression of AKAP95. The numbers of exon junction reads and PSI are indicated. P values by two-sided Student’s *t*-test for a and b. Uncropped blots are provided as source data.

**Extended Data Figure 3.**
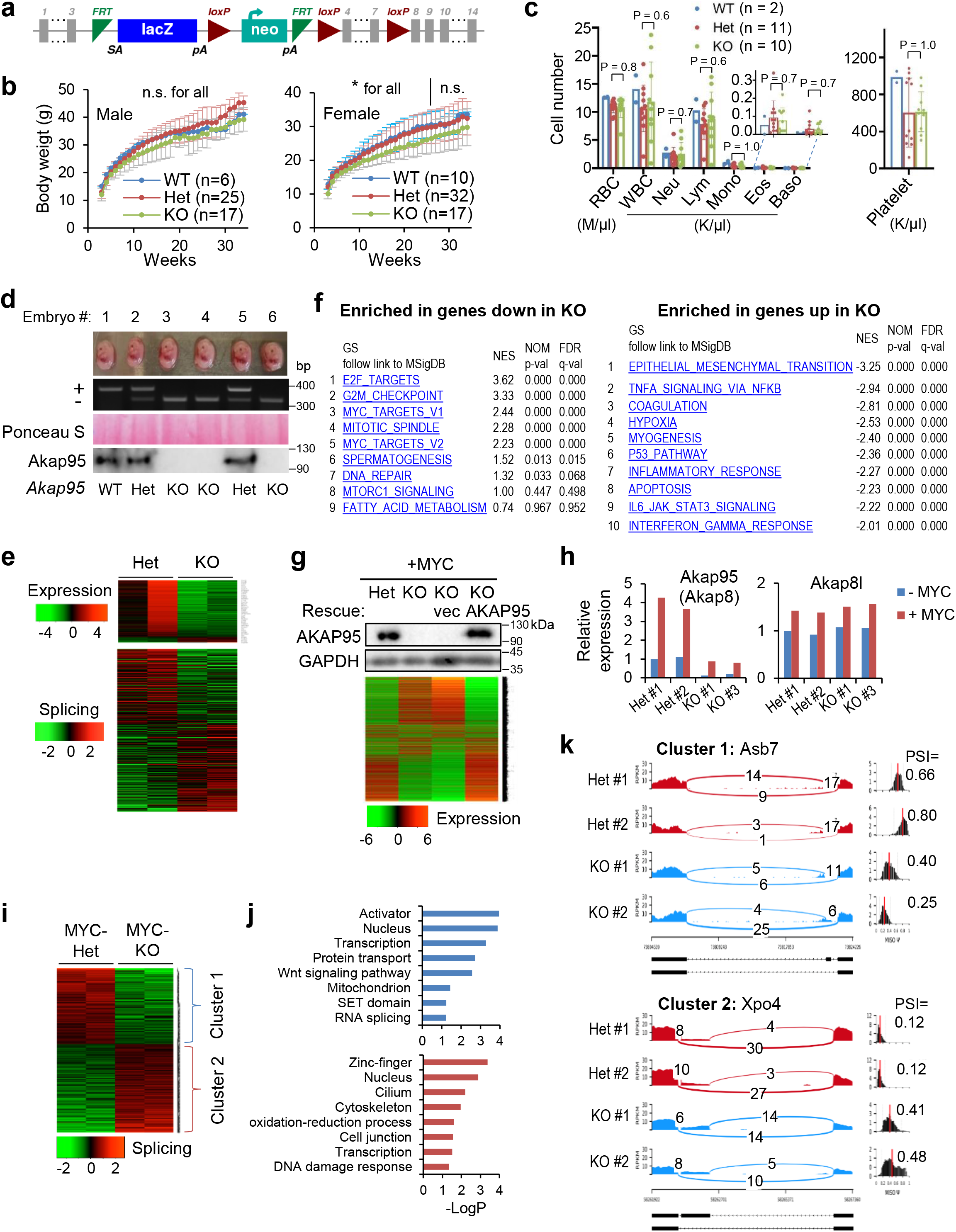
AKAP95 is required for transformation and overcoming oncogene-induced cellular senescence. **a,** Schematic of the “Knockout-first” *Akap95* (*Akap8*) allele (null without further recombination). **b,** Body weights of mice with indicated *Akap95* genotypes. Not significant (n.s.) between any two groups at any time for male, or after week 26 for female. *between Het and KO before week 26 for female. **c,** Peripheral blood profiles of 8-week mice of indicated genotype. **d,** MEFs were derived from mouse embryos. Images of six embryos from the same litter were shown at the top, followed with genotyping results, Ponceau S staining, and immunoblotting of MEFs. **e,** Top, heatmap showing relative expression levels of genes and clustered by changes in un-transduced KO MEFs from 2 embryos each. It includes 203 and 20 genes down- or up-regulated in KO, respectively. Bottom, heatmap showing relative alternative splicing of genes clustered by PSI changes. It includes 285 and 332 alternative splicing events with decreased or increased PSI in KO, respectively. Also see Supplementary Table 2, tab 6 and 7. **f,** Top 10 gene sets enriched in genes down-(left, n = 265) and up-(right, n = 742) regulated in the MYC-transduced KO versus Het MEFs. **g,** Rescue of the gene expression profile by introduction of human AKAP95 into the MYC-transduced KO MEFs as shown by immunoblotting and heatmap for relative expression of down- or up-regulated genes in the indicated cells. Also see Supplementary Table 2, tab 2. Repeated 2 times. **h,** Relative expression of *Akap95* and *Akap8l* in un-transduced (-MYC) and MYC-transduced (+MYC) MEFs from n = 2 Het and two KO embryos, as determined by normalized RNA-seq reads. **i,** Heatmap showing relative alternative splicing of genes clustered by PSI changes in MYC-transduced MEFs from 2 embryos each. It includes 216 and 252 alternative splicing events with decreased or increased PSI in KO versus Het MEFs, respectively. Also see Supplementary Table 2, tab 3. **j,** Gene ontology analysis for the indicated gene clusters from the heatmap in **i**. Blue (n = 216) and red (n = 252) show functions significantly enriched in genes with PSI increase or decrease by KO, respectively. **k,** Sashimi plots showing alternative splicing changes for each gene cluster from the heatmap using two examples, Asb7 for cluster 1, and Xpo4 for cluster 2. ns, not significant, *P < 0.05, by two-sided Student’s *t*-test for b, one-way ANOVA for c and modified Fisher’s exact test for f, j. Uncropped blots are provided as source data.

**Extended Data Figure 4.**
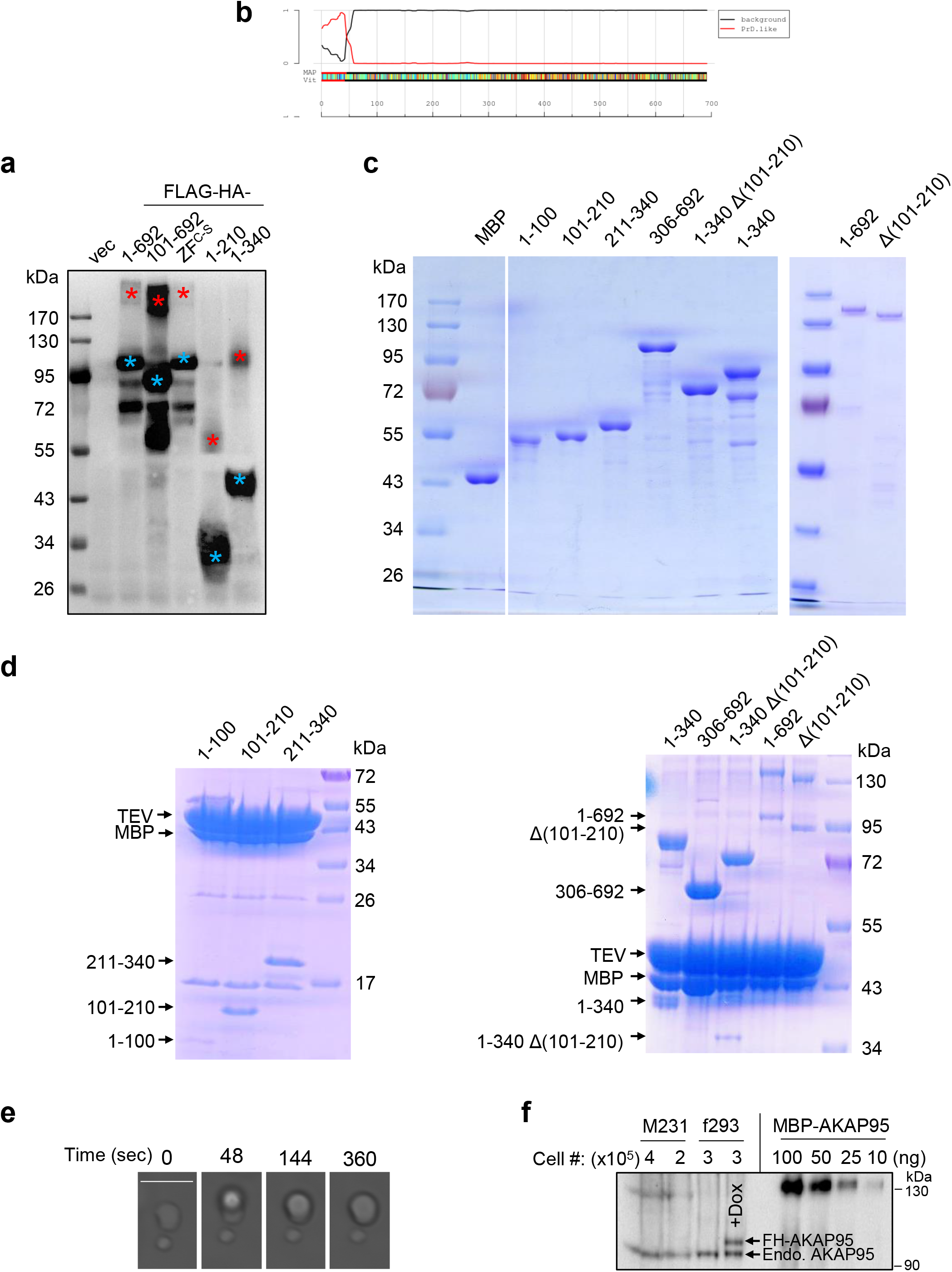
AKAP95 phase separation in vitro. **a,** 293T cells were transfected with either empty vector (vec), or indicated AKAP95 construct with FLAG-HA-tag. Following α-Flag IP, the pulldown proteins were boiled and resolved by SDS-PAGE and detected by immunoblotting with α-HA. Blue and red asterisks indicate monomer and dimer, respectively. **b,** Identification of 1-100 as a probable prion subsequence on AKAP95. By the PLAAC program, using homo sapiens as background and core length of 30. **c,** Purified MBP and MBP fused to AKAP95 truncations as indicated or full-length AKAP95 (1-692) were resolved on SDS-PAGE and stained with Coomassie blue. **d,** MBP fused to AKAP95 truncations as indicated or full-length AKAP95 were resolved on SDS-PAGE and stained with Coomassie blue following treatment with TEV protease. Note that the cleaved MBP serves as a better indicator for cleavage efficiency as staining signal various for protein fragments of different sequences and sizes. **e,** Another event of fusion of two droplets formed by 50 μM MBP-AKAP95 (101-210) in 30 mM NaCl and 10% of PEG6000 after treatment with TEV protease for 30 min. Scale bar, 5 μm. Also see Movie S1. **f,** Quantification of nuclear AKAP95 concentration by anti-AKAP95 Western blot. Total lysates from indicated number of MDA-MB-231 (M231) and flp-TREx 293 cells (f293, un-induced and dox-induced for FH-AKAP95 expression) were loaded, along with indicated ng of purified MBP-AKAP95. AKAP95 signal of un-induced f293 is similar to that of 25 ng of MBP-AKAP95. All experiments were repeated 2 times. Uncropped blots are provided as source data.

**Extended Data Figure 5.**
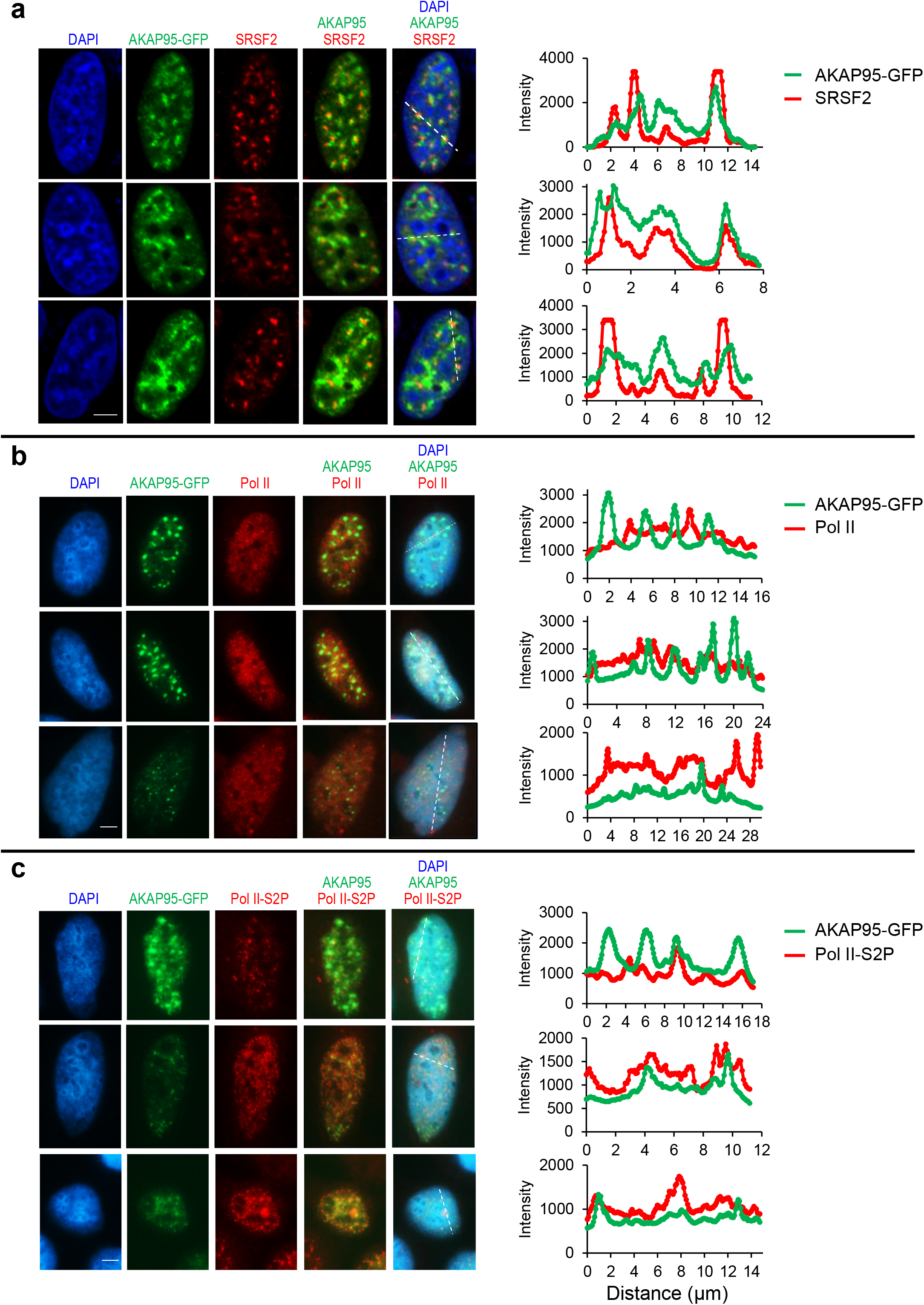
AKAP95 partially localizes in nuclear speckles and with actively transcribing Pol II. Fluorescence microscopy images of HeLa cells transiently expressing AKAP95-GFP. Nucleus DNA was stained by DAPI, and specific proteins were stained with antibodies for SRSF2 (**a**), Pol II (**b**), and Pol II-S2P (**c**). At least 10 cells were imaged for each staining and show similar trend. Right, quantification of the signal intensity of indicated molecules across the dotted lines shown in the images. Quantification by Image J. Scale bar, 5 μm for all.

**Extended Data Figure 6.**
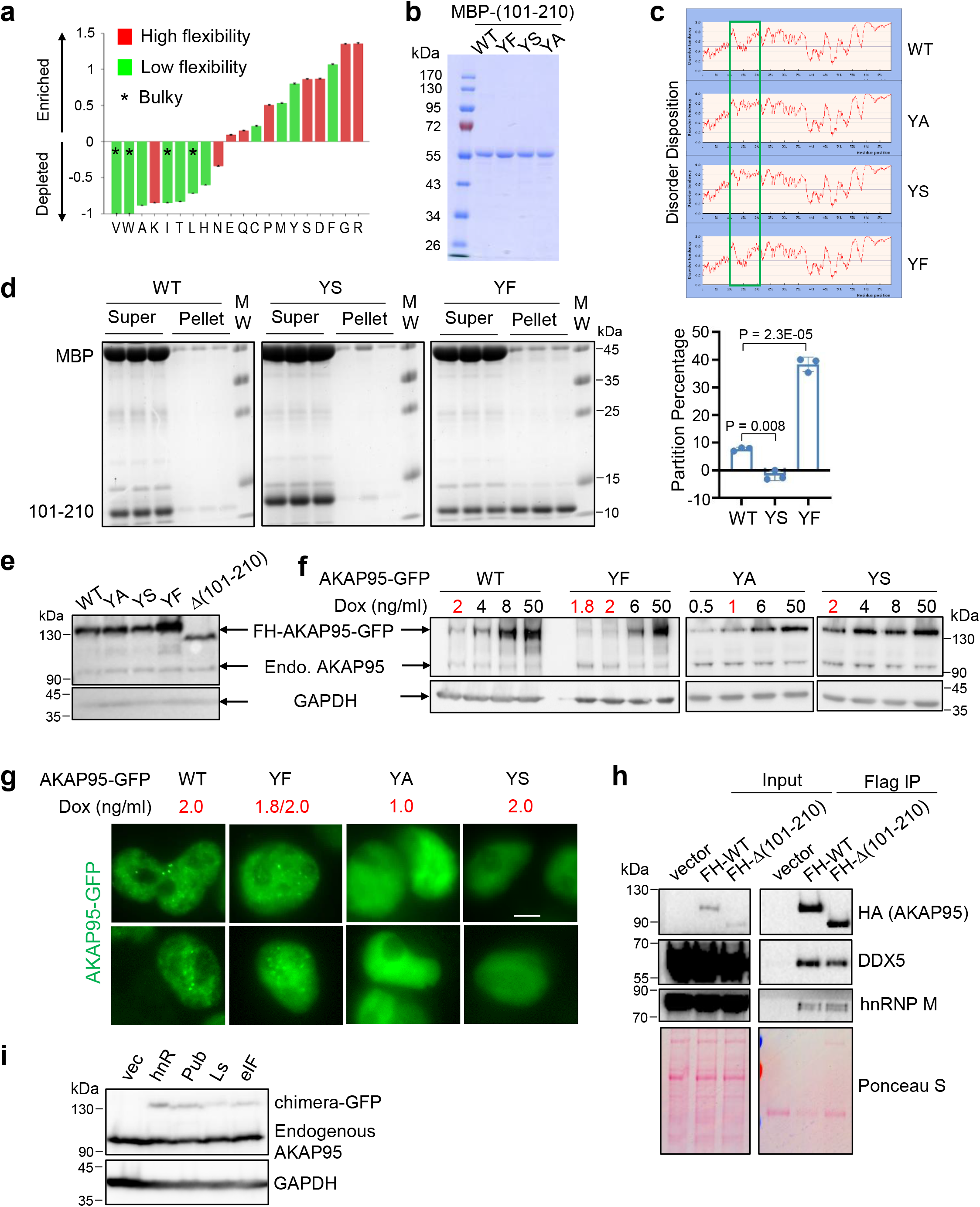
AKAP95 condensation requires Tyrosine in 101-210 and regulates splicing. **a,** Amino acid enrichment for AKAP95 (101-210). By Composition Profiler, using SwissProt 51 Dataset as background. **b,** Purified MBP fused to AKAP95 (101-210) WT and mutants were resolved on SDS-PAGE and stained with Coomassie blue. Repeated 2 times. **c,** Disorder plot of AKAP95 WT or mutants with indicated mutations in 101-210. **d,** MBP-AKAP95 (101-210) WT, YS, and YF, all at 35 μM and in 30 mM NaCl, were treated with TEV protease for 2 hrs in 3 independent assays, and subjected to centrifugation. The supernatant and pellets (resuspended in the same volume as the supernatant) were resolved by SDS-PAGE followed with coomassie blue staining. MBP signal in the pellet reflects residual supernatant fraction, and its percentage [MBP pellet/(supernatant + pellet)] was subtracted from the (101-210) pellet percentage. Such normalized (101-210) pellet percentages are plotted as Partition Percentage as mean ± SD of n = 3 independent experiments. It is most likely that all supernatants may also have substantial portion of condensates. Moreover, the size cutoff of condensates is also arbitrary, as protein assemblies may take a continuum of size distribution^2^. **e,** Immunoblotting by α-AKAP95 (top) or GAPDH (bottom) of total lysates from Flp-In T-Rex 293 cell lines induced to express full-length AKAP95 WT or indicated mutants fused to GFP. Repeated > 2 times. **f,g,** Indicated full-length AKAP95 WT or mutants fused to GFP were induced by various concentrations of doxycycline in Flp-In T-Rex 293 cell lines. Immunoblotting of total cell lysates with indicated antibodies (**f**). The doxycycline concentrations in red font activated the transgene at the near endogenous level, and were selected for treating cells and fluorescence microscopy assays of fixed cells in (**g**). Scale bar, 5 μm. Repeated 2 times. **h,** 293T cells transiently expressing indicated constructs with FLAG-HA-tag were used for α-FLAG immunoprecipitation and immunoblotting with indicated antibodies and Ponceau S staining. Repeated 2 times. **i,** Immunoblotting of 293T cells transfected with empty vector or indicated FLAG-HA-tagged AKAP95 chimeras fused to GFP. Bottom, by anti-GAPDH. Top, by anti-AKAP95 (Bethyl Laboratories, A301-062A, recognizes an epitope in a region between residue 575 and 625 of human AKAP95). Repeated 2 times. P values by two-sided Student’s *t*-test for d. Uncropped blots are provided as source data.

**Extended Data Figure 7.**
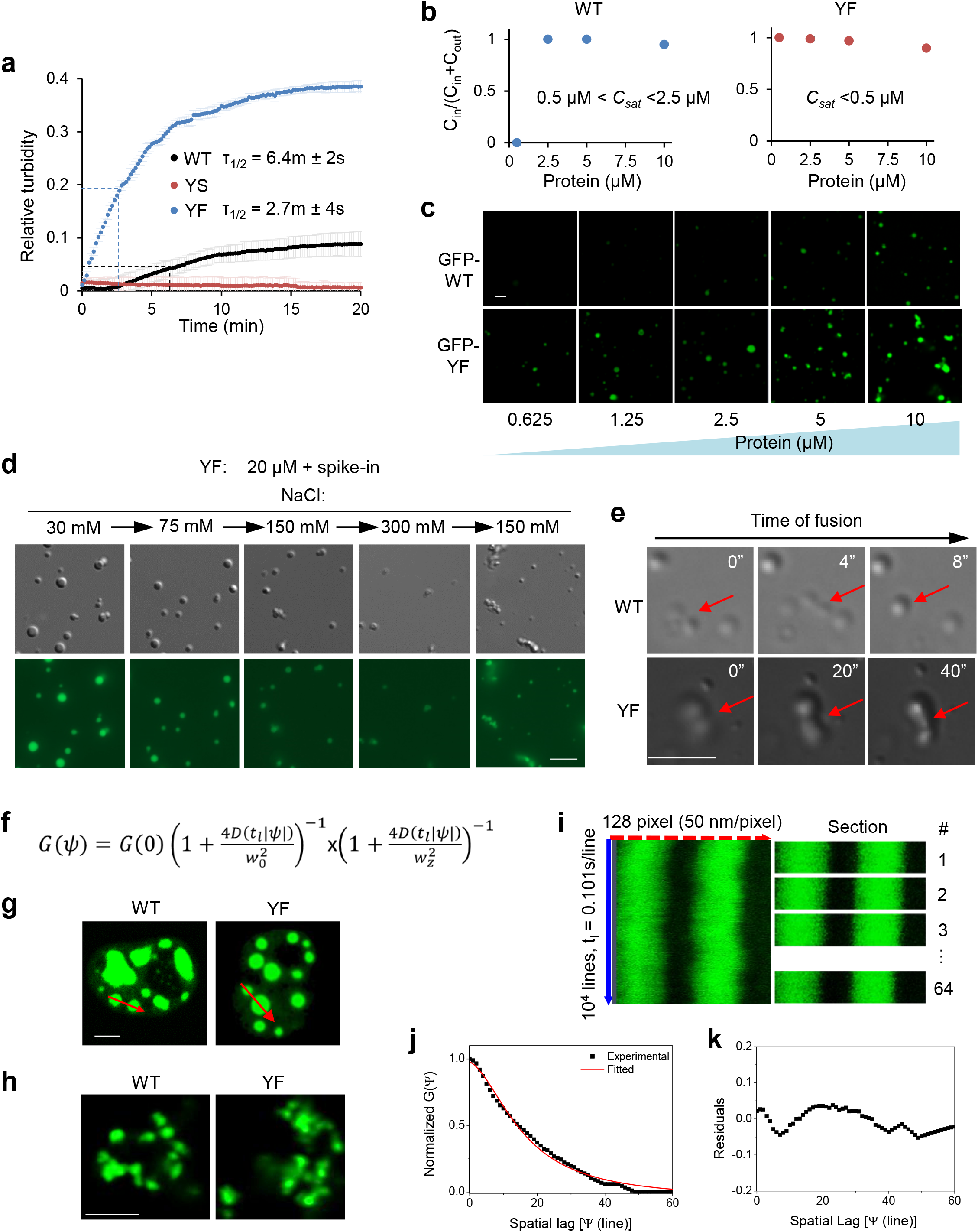
YF mutation alters material properties of AKAP95 condensates. **a,** OD450 at different time after TEV protease treatment of 50 μM MBP-AKAP95 (101-210) WT, YS, and YF in 150 mM NaCl, as mean ± SD of readings after subtracting that of MBP at each time (constant at 0.07-0.08) from n = 3 independent assays. Dashed lines show half of the maximum turbidity and time (τ_1/2_) to reach it. **b,** Ratio of protein concentration inside the droplets over sum of inside and outside for (101-210) WT and YF at increasing protein concentrations and in 30 mM NaCl, as mean ± SD from n = 6 randomly picked droplets each based on Fig. 7a. **c,** Confocal microscopy images of GFP-AKAP95 (101-210) WT and YF at increasing protein concentrations, all in 150 mM NaCl and 10% of PEG6000 after TEV protease treatment for 20 min. **d,** DIC and fluorescence microscopy images for 20 μM MBP-AKAP95 (101-210) YF spiked with Oregon-green-labeled same protein (molar ratio 10:1), after TEV protease treatment for 30 min. Changes in NaCl concentration is indicated. Images were taken 5 min after salt adjustment. **e,** Different extent of droplet fusion (arrows) by AKAP95 (101-210) WT and YF, both at 50 μM and in 30 mM NaCl and 10% of PEG6000 after TEV protease treatment for 30 min. A similar trend was observed in > 5 fusion events (or attempted fusion for YF) for each. Also see Movies 4-6. **f,** Equation for Line Raster Scan Image Correlation fitting autocorrelation G(Ψ), which depends on G(0)=γ/N (γ: beam profile, N = number of mobile particles), diffusion coefficient D (μm^2^/s), line scan time t_l_, pixel size Ψ, radial beam waist w_0_ and axial beam waist w_z_. The radial waist w_0_ (0.218 μm) was calibrated with sub-diffraction beads (0.1 μm) diluted solution as reported before^3^. The axial waist w_z_ was considered equal to 3*w_0_. **g,** Fluorescence confocal microscope image of HeLa cell expressing full-length AKAP95 WT or YF fused to GFP. Arrow indicates the scanned region for in vivo Line RICS. **h,** Fluorescence confocal microscope image of GFP-AKAP95 (101-210) WT and YF in 150 mM NaCl and 10% of PEG6000 after TEV protease treatment for 20 min, for in vitro Line RICS experiments. **i,** Line Fluorescence carpet formed by ~10^4^ lines (each line is composed by 128 pixels, 50 nm/pixel) acquired with 0.101 s line scan time and 32.8 μs/pixel. Line RICS autocorrelation curves were computed on 64 sections of 128 lines followed by averaging all the curves. The sectioning of the line carpet permitted to avoid the effect of the movement of the condensates on the measurement of the autocorrelation curves. (λex=488 nm). **j,** Line RICS autocorrelation curves for experimental data and fitted. **k,** Residuals of the fitting. All experiments were Repeated > = 2 times. Scale bar, 2 μm for c and h, 5 μm for d, e, and g.

**Extended Data Figure 8.**
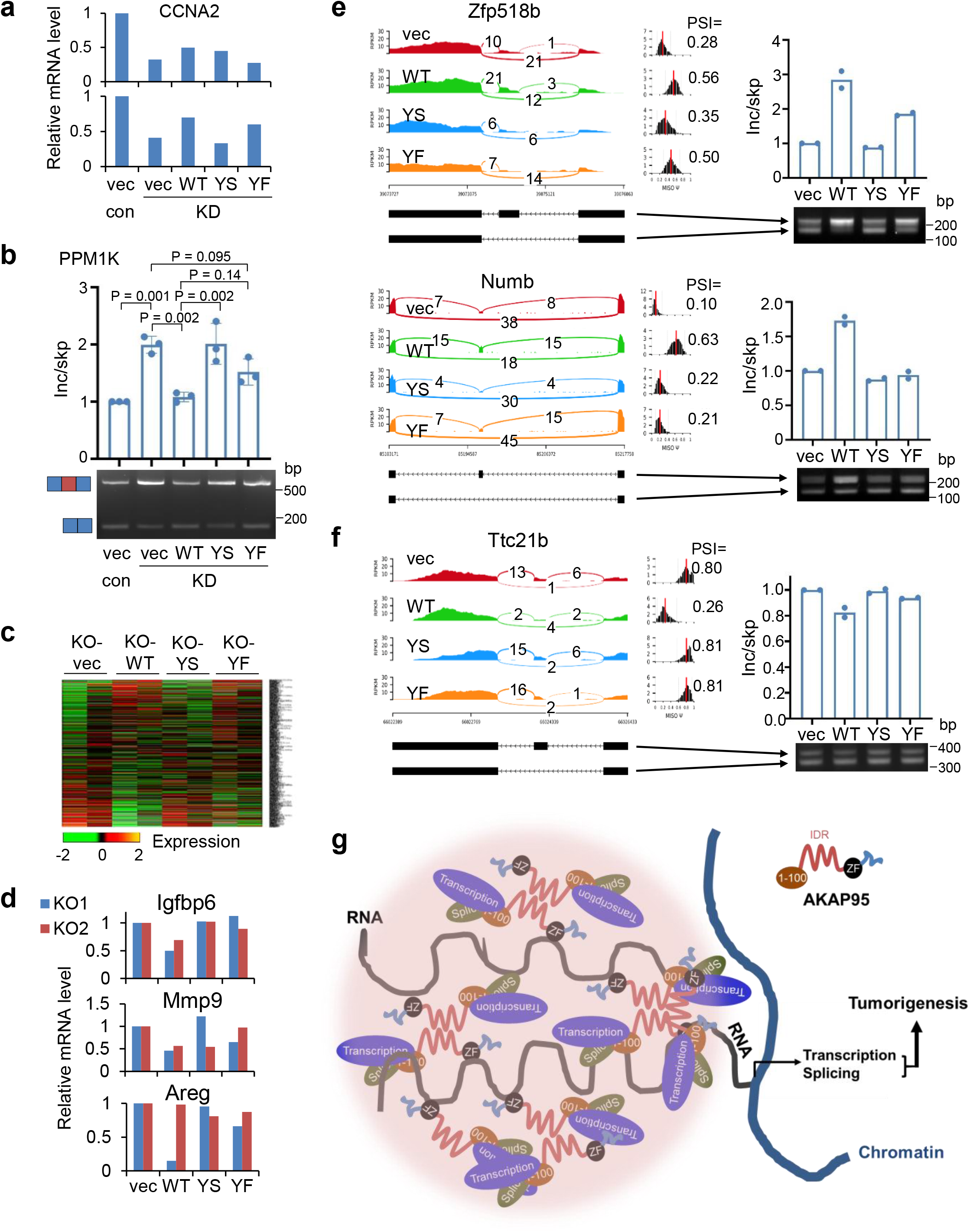
Regulation of tumorigenesis and gene expression by AKAP95 requires its condensation with appropriate material properties. **a,b,** MDA-MB-231 cells were virally infected to stably express scramble (control) or AKAP95 shRNA #1 (KD) and the indicated constructs including empty vector (vec) and FLAG-HA-tagged full-length AKAP95 WT or mutants. **a,** Relative *CCNA2* mRNA level as determined by RT-qPCR and normalized to *GAPDH*, and plotted for each of the 2 biological repeats individually. **b,** RT-PCR for ratios for exon-included over-skipped *PPM1K* transcript, as mean ± SD from n = 3 biological repeats. **c-f,** MYC-transduced *Akap95* KO MEFs were transduced with vector or constructs expressing HA-tagged full-length AKAP95 WT or mutants. **c,** Heatmap showing relative expression levels of genes changed in MYC-transduced KO MEFs (from 2 embryos each) stably expressing indicated rescue constructs. Also see Supplementary Table 2, tab 4. **d,** Relative mRNA levels of indicated SASP genes as determined by RNA-seq reads from n = 2 biological repeats (KO1 and KO2). **e,f,** Sashimi plots showing example genes for which the alternative exon inclusion was promoted (**e**) or suppressed (**f**) by introduction of AKAP95 WT, but not as effectively by YS or YF, and RT-PCR for the inclusion of the alternative exon, as mean ± SD from n = 2 embryos each. **g,** A model for how AKAP95 condensates may regulate gene expression for tumorigenesis. P values by one-way ANOVA followed by Tukey’s post hoc test. Uncropped blots are provided as source data.

### Supplementary Table Legends

**Supplementary Table 1. Changes of gene expression and alternative splicing by AKAP95 KD and rescue in TNBC cells**

This table contains 2 tabs as explained below:

**Tab 1. Gene expression changes by AKAP95 KD and rescue in TNBC cells.**

By expression analyses of the RNA-seq results from MDA-MB-231 cells that were virally infected to stably express scramble control shRNA or AKAP95 shRNA (KD) as well as vector or AKAP95 construct. Shown are steps to transform the expression values to generate heatmap as described in Materials and Methods.

**Tab 2. Changes of gene alternative splicing by AKAP95 KD and rescue in TNBC cells.**

By splicing analyses of the RNA-seq results from MDA-MB-231 cells that were virally infected to stably express scramble control shRNA or AKAP95 shRNA (KD) as well as vector or AKAP95 construct.

**Supplementary Table 2. Changes of gene expression and alternative splicing by** *Akap95* **KO and rescue in MYC-transduced MEFs**

This table contains 7 tabs as explained below:

**Tab 1. Gene expression changes by** *Akap95* **KO in MYC-transduced MEFs.**

By expression analyses of the RNA-seq results from MYC-transduced *Akap95* Het and KO MEFs derived from n = 2 embryos for each. Shown are steps to transform the expression values to generate heatmap as described in Materials and Methods.

**Tab 2. Gene expression changes by** *Akap95* **KO and rescue in MYC-transduced MEFs.**

By expression analyses of the RNA-seq results from MYC-transduced *Akap95* Het, and KO MEFs, and KO MEFs that were introduced with vector or AKAP95 WT.

**Tab 3. Changes of gene alternative splicing by** *Akap95* **KO in MYC-transduced MEFs.**

By splicing analyses of the RNA-seq results from MYC-transduced *Akap95* Het and KO MEFs derived from n = 2 embryos for each.

**Tab 4. Gene expression changes by AKAP95 expression in MYC-transduced** *Akap95* **KO MEFs.** By expression analyses of the RNA-seq results from MYC-transduced *Akap95* KO MEFs (from n = 2 embryos) that were introduced with vector, AKAP95 WT, YS, or YF mutant.

**Tab 5. Changes of gene alternative splicing by AKAP95 expression in MYC-transduced** *Akap95* **KO MEFs.**

By splicing analyses of the RNA-seq results from MYC-transduced *Akap95* KO MEFs (from n = 2 embryos) that were introduced with vector, AKAP95 WT, YS, or YF mutant.

**Tab 6. Gene expression changes by AKAP95 expression in** *Akap95* **KO MEFs.**

By expression analyses of the RNA-seq results from un-transduced *Akap95* Het and KO MEFs derived from n = 2 embryos for each.

**Tab 7. Changes of gene alternative splicing by AKAP95 expression in** *Akap95* **KO MEFs.**

By splicing analyses of the RNA-seq results from un-transduced *Akap95* Het and KO MEFs derived from n = 2 embryos for each.

**Supplementary Table 3. AKAP95 chimeras information**

Information of the IDRs from 4 different proteins that were used to replace the 101-210 region of AKAP95 to make AKAP95 chimeras.

**Supplementary Table 4. Information for Primers, shRNAs, and siRNAs**

### Supplementary Movie Legends

**Supplementary Movie 1. AKAP95 droplets in vitro are highly dynamic and can fuse.**

Purified MBP-AKAP95 (101-210) formed droplets at 50 μM, in 30 mM NaCl and 10% of PEG6000 after treatment with TEV protease for 30 min. The yellow triangle points to a fusion event of two droplets. Played at 134x speed. Repeated > = 3 times.

**Supplementary Movie 2. Rapid fusion of AKAP95 foci in the cell nucleus.**

Full-length AKAP95 (ZF^C-S^)-GFP foci in a HeLa cell nucleus following transfection. The arrows point to two different fusion events. Played at 20x speed. Repeated > = 3 times.

**Supplementary Movie 3. FRAP assays for intracellular AKAP95.**

Representative FRAP assays for AKAP95 fused to GFP and transfected into HeLa cells. Played at 20x speed. Repeated > = 3 times.

**Supplementary Movie 4. Different extent of droplet fusion by AKAP95 WT and YF.**

Purified AKAP95 (101-210) WT and YF both at 50 μM in 30 mM NaCl and 10% of PEG6000 after treatment with TEV protease for 30 min. Yellow square and green circle highlight the fusion event for YF and WT, respectively. These movies are the first 100 seconds excerpts from Supplementary Movies 5 and 6. Repeated > = 3 times.

**Supplementary Movies 5 and 6. AKAP95 WT and YF droplets in vitro.**

Purified AKAP95 (101-210) WT (Movie 5) and YF (Movie 6) both at 50 μM in 30 mM NaCl and 10% of PEG6000 after treatment with TEV protease for 30 min, and were videoed under microscope for 60 min in total, with one image every 4 seconds. Played at 20x speed. Repeated > = 3 times.

## Notes

### Competing Interest Statement

The authors have declared no competing interest.

### Summary of Updates

This revision has added experiments to show altered diffusion by the YF mutation, and added authors from UC Irvine, as well as re-reformatting of the paper.

